# Receptor trafficking couples intracellular auxin perception to rapid signaling

**DOI:** 10.64898/2026.04.10.716253

**Authors:** Jing Wang, Jiang-ping Ye, Minmin Zhang, Hanqian Feng, Mengzhen Liu, Yuanzhi Huang, Yongqiang Yu, Tongda Xu, Baiyan Lu, Chao Li

## Abstract

Rapid auxin responses are triggered by Auxin-Binding Protein 1 (ABP1) signaling; however, the coupling of hormone perception to signaling activation is unclear. ABP1, a cell surface auxin receptor, is localized to endoplasmic reticulum (ER), raising a fundamental question of where auxin is perceived and how signaling-competent receptors are mobilized to cell surface. Herein, we established a dark-induced auxin synthesis system in *Arabidopsis* hypocotyl; *abp1* mutants showed pronounced elongation. Auxin dose-dependently stimulates LORELEI-like glycosyl-phosphatidylinositol-anchored protein 1 (GPI-AP, LLG1) interaction with ABP1, driving their outward trafficking from ER. However, this complex dissociated in acidic apoplast, facilitating H^+^-ATPase-mediated auxin signaling. Our findings reveal how intracellular auxin perception is converted into rapid cell-surface signaling, establishing a mechanistic framework for fast auxin responses and uncovering a GPI-AP-mediated trafficking mechanism for ER-retained receptors.

## Introduction

Rapid cellular responses depend on mechanisms that couple hormone perception with immediate signal activation. In plants, auxin centrally regulates growth and development (*1*); beyond its well-known transcriptional effects (*2*), it induces rapid responses through cell-surface signaling pathways, such as proton extrusion, membrane depolarization, calcium transients, and cytoplasmic streaming (*3-6*). This pathway operates independently of transcriptional regulation, controlling diverse developmental processes, including cell expansion and tropical growth (*7, 8*). Auxin-binding protein 1 (ABP1)- and transmembrane kinase receptor-like kinases (TMKs)-mediated pathways have recently been established (*9, 10*).

ABP1 has long been proposed as an auxin receptor (*11*). Isolated originally from maize owing to its auxin-binding activity, ABP1 was recently established as a key component of the cell-surface auxin signaling pathway (*12, 13*). ABP1 forms an auxin-dependent receptor complex with TMKs, triggering an ultrafast global phospho-signaling cascade that activates plasma membrane (PM) H^+^-ATPase and promotes cell wall acidification and vascular regeneration (*9, 10*). ABP1-like proteins participate in TMK-mediated signaling and regulate developmental processes, such as pavement cell interdigitation and hypocotyl elongation, suggesting partially overlapping but distinct functions within the ABP1/ABLs signaling module (*14*). Although ABP1 has emerged as a central component of rapid auxin signaling, the physiological significance of this pathway is unclear because genetic disruption of ABP1 often results in unexpectedly mild phenotypes (*15, 16*).

Moreover, ABP1’s subcellular localization raises a long-standing paradox (*11*). Although ABP1 functions in the cell surface signaling pathway, it contains a KDEL motif that conserved in angiosperms (*17*). Therefore, ABP1 is predominantly retained within the ER under steady-state conditions (*18*). Auxin treatment promotes ABP1 secretion from the ER into the apoplast in both *Arabidopsis* and *maize* (*10, 19*), suggesting that receptor mobilization is linked to auxin signaling. Biochemical studies have shown preferential binding of ABP1 to auxin under acidic conditions (*10, 20*), leading to the prevailing view that auxin perception occurs at the cell surface. However, this model does not explain how auxin triggers ABP1 export from the ER. Thus, two fundamental questions remain unresolved: whether ABP1 directly senses auxin within the ER, and how auxin perception overcomes KDEL-mediated ER retention to mobilize ABP1 from intracellular compartments to the cell surface, thereby initiating rapid signaling responses (*11, 21*). Here, we resolve how an ER-retained auxin receptor mediates rapid PM signaling by uncovering an auxin-dependent ABP1–LLG1 trafficking pathway that generates signaling-competent receptors at the cell surface.

## Results

### Dark treatment induces ABP1 trafficking through auxin biosynthesis for hypocotyl elongation

In *pABP1:ABP1-eGFP* transgenic plants, ABP1 expression in the hypocotyl epidermis and cortex exhibited temporal variation, peaking between days 4 and 5 (fig. S1A–B). Notably, treatment of 4-day-old seedlings to the dark induced significantly longer hypocotyls in *abp1-c1* and *abp1-TD1* mutants than in the wild type (WT) (Fig. 1A). Growth kinetics analysis using a dual-point tracking assay (*22*) revealed that hypocotyl elongation proceeded through two clearly distinguishable phases: an initial fast-growing phase followed by a slower, sustained growth phase. Throughout the growth period, both *abp1* mutants exhibited significantly increased newly formed growth with higher rates of hypocotyl growth compared to the WT. Specifically, *abp1-c1* maintained accelerated growth throughout the monitoring period, whereas *abp1-TD1* displayed rapid elongation only during the early phase, followed by a gradually declining growth rate during later stages. Ultimately, both mutants developed longer hypocotyls than the WT, with the elongation phenotype being more pronounced in *abp1-c1* (Fig. 1B). The upper hypocotyl region was used for all cytological analyses. Notably, cortical cell size significantly increased in *abp1* mutants after dark treatment (Fig. 1C and fig. S1C–D). Under standard long-day conditions, *ABP1* predominantly colocalized with the ER marker RFP-HDEL rather than with FM4-64-stained PM, indicating its ER location (*23*) (fig. S2A). In contrast, dark treatment promoted a marked redistribution of ABP1 toward the PM. Consistently, Pearson’s correlation coefficient analysis confirmed a significant increase in ABP1-FM4-64 colocalization after exposure to the dark compared to control conditions, suggesting a time-dependent re-localization of ABP1 from the ER to the PM (fig. S2B–C). Thus, the ABP1-regulated hypocotyl elongation process helped us establish an effective, tractable system to elucidate the molecular mechanisms underlying ABP1 function.

**Figure 1.**
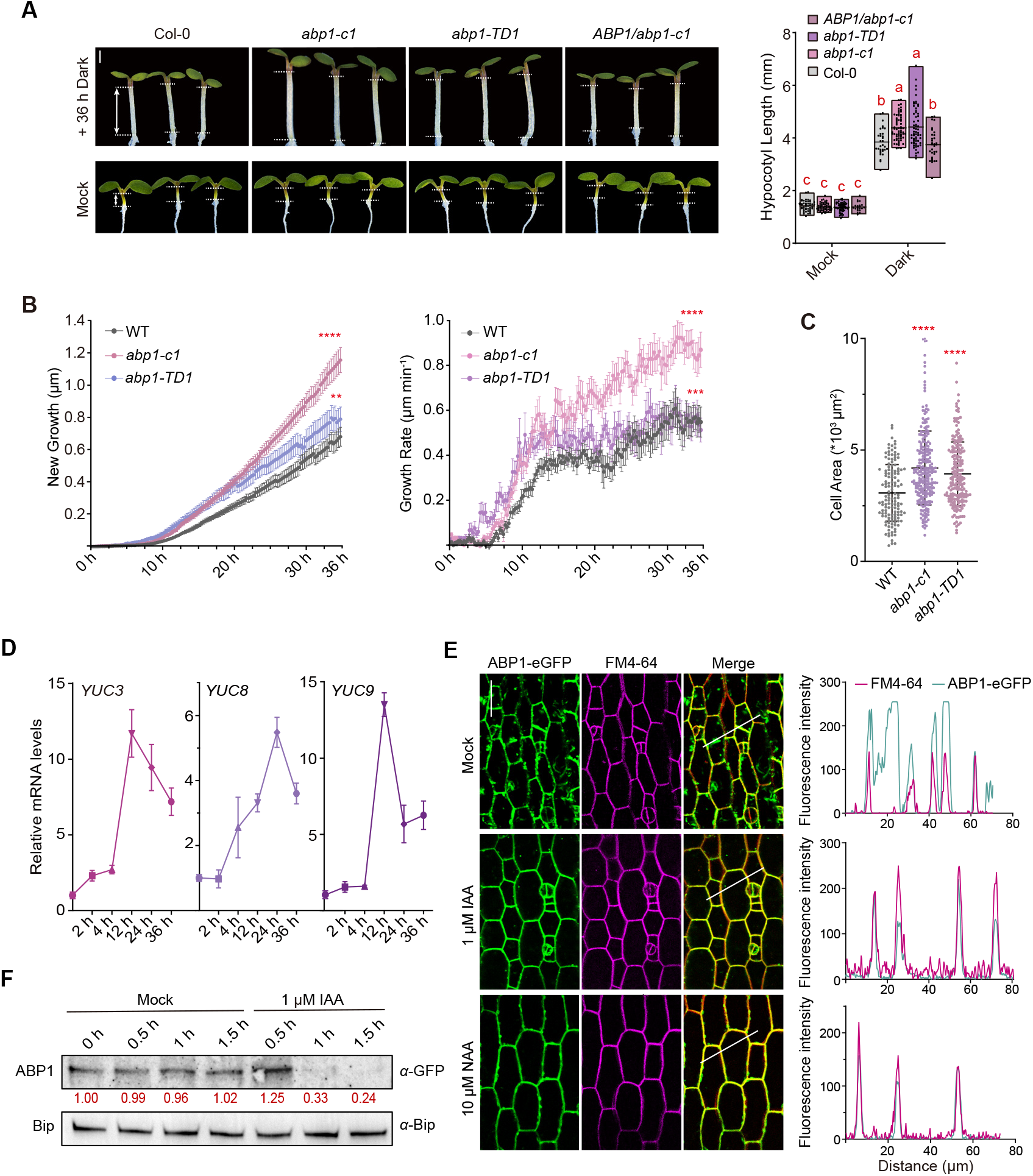
Dark treatment induces ABP1 trafficking through auxin biosynthesis for hypocotyl elongation. (A) Phenotype analysis and quantification of hypocotyl length in 4-day-old Col-0, *abp1-c1, abp1-TD1 and pABP1:ABP1-eGFP/abp1-c1* seedlings subjected to mock or dark treatment for 36 h. *n* ≥ 21 hypocotyls. Scale bar, 2 mm. (B) Time-lapse analysis of hypocotyl new growth and growth rate in 4-day-old WT, *abp1-c1* and *abp1-TD1* seedlings during continuous dark treatment for 36 h. Images were captured every 15 min. *n* ≥ 28 hypocotyls. (C) Quantification of hypocotyl cell areas in 4-day-old WT, *abp1-c1* and *abp1-TD1* seedlings subjected to dark treatment for 36 h. *n* ≥ 135 cells from 24–39 hypocotyls. (D) RT-qPCR analysis of *YUC3, YUC8* and *YUC9* in aboveground part of 4-day-old seedlings subjected to darkness for 0, 2, 4, 12, 24 or 36 h. Transcript levels were expressed as fold change relative to 0 h (set to 1). (E) Confocal microscopy images (left) and fluorescence plot profiles (right) of hypocotyl epidermal cells in 4-day-old seedlings treated with/without IAA or NAA for 1 h. FM4-64 labels the PM. White lines indicate fluorescence measurement regions. Scale bar, 20 µm. (F) Western blot analysis of ABP1-eGFP protein levels following IAA treatment in ER fraction. α-Bip serves as ER marker. Red numbers indicate relative ABP1-eGFP protein levels at each time point relative to untreated control. In (A–C), data are presented as mean ± SD. Significant differences were determined using two-way ANOVA (A). Different lowercase letters indicate significant differences at *P* < 0.05. Significant differences were determined using Student’s *t*-test (B, C). \*\**P* < 0.01, \*\*\**P* < 0.001, \*\*\*\**P* < 0.0001.

Consistent with previous reports that darkness promotes auxin biosynthesis in cotyledons (*24*), free IAA levels increased markedly after 24 h of dark treatment in the aboveground seedling tissues (fig. S3A). This increase was accompanied by the induction of the expression of auxin biosynthetic genes (*25*), including *ASA1, TAA1*, and *YUC3/8/9*, and enhanced *pYUC9* activity in hypocotyls (Fig. 1D and fig. S3B–D). Thus, darkness-induced auxin accumulation inhibited hypocotyl elongation (*26*). Consistently, the auxin synthesis inhibitor, PPBo, significantly suppressed darkness-induced hypocotyl elongation. Notably, the pronounced differences between the WT and *abp1* mutants under dark conditions were largely abolished by PPBo treatment (fig. S3E). Thus, in the context of darkness treatment, the overaccumulation of auxin leads to an ABP1-dependent pathway that inhibit hypocotyl elongation (*26*).

Next, we tested whether increased auxin levels promoted ABP1 trafficking. Auxin treatment rapidly increased the colocalization of ABP1-eGFP with the plasma membrane marker FM4-64 and promoted its redistribution from the intracellular compartments to the plasma membrane (Fig. 1E and fig. S4A–B). Consistently, ABP1 abundance decreased in the ER fraction without changes in the total protein levels (Fig. 1F and fig. S4C). Conversely, ABP1 was largely localized intracellularly in the low-auxin *asa1* mutant (fig. S4D). These results suggest that darkness-induced auxin accumulation drives ABP1 trafficking from the ER to the PM.

### Dark-induced LLG1 expression facilitates ABP1 trafficking through protein-protein interaction

Given the functional studies on the LRE family glycosylphosphatidylinositol-anchored proteins (GPI-APs) as chaperones that mediate protein trafficking and the involvement of LLG1 in auxin signaling (*27-29*), we next investigated whether LRE/LLGs were involved in ABP1 secretion. Among vegetative tissues that expressed LRE and LLG1 (*30*), LLG1 showed a drastically higher expression in the hypocotyls (fig. S5A). In the hypocotyl epidermis and cortex of the *pLLG1:eGFP-LLG1* transgenic seedlings, *eGFP-LLG1* fluorescence peaked around days 3–4, following a temporal pattern similar to *ABP1* (fig. S5B). Similar to *abp1* mutants, the *llg1-2* mutant showed dark-induced accelerated hypocotyl elongation compared with the WT (fig. S5C). Notably, in a lanolin-paste auxin application assay using cotyledons (*31*), we observed longer hypocotyls in *abp1-c1, abp1-TD1, llg1-2*, and *abp1-c1 llg1-2* mutants compared with the WT (Fig. 2A and fig. S5D).

**Figure 2.**
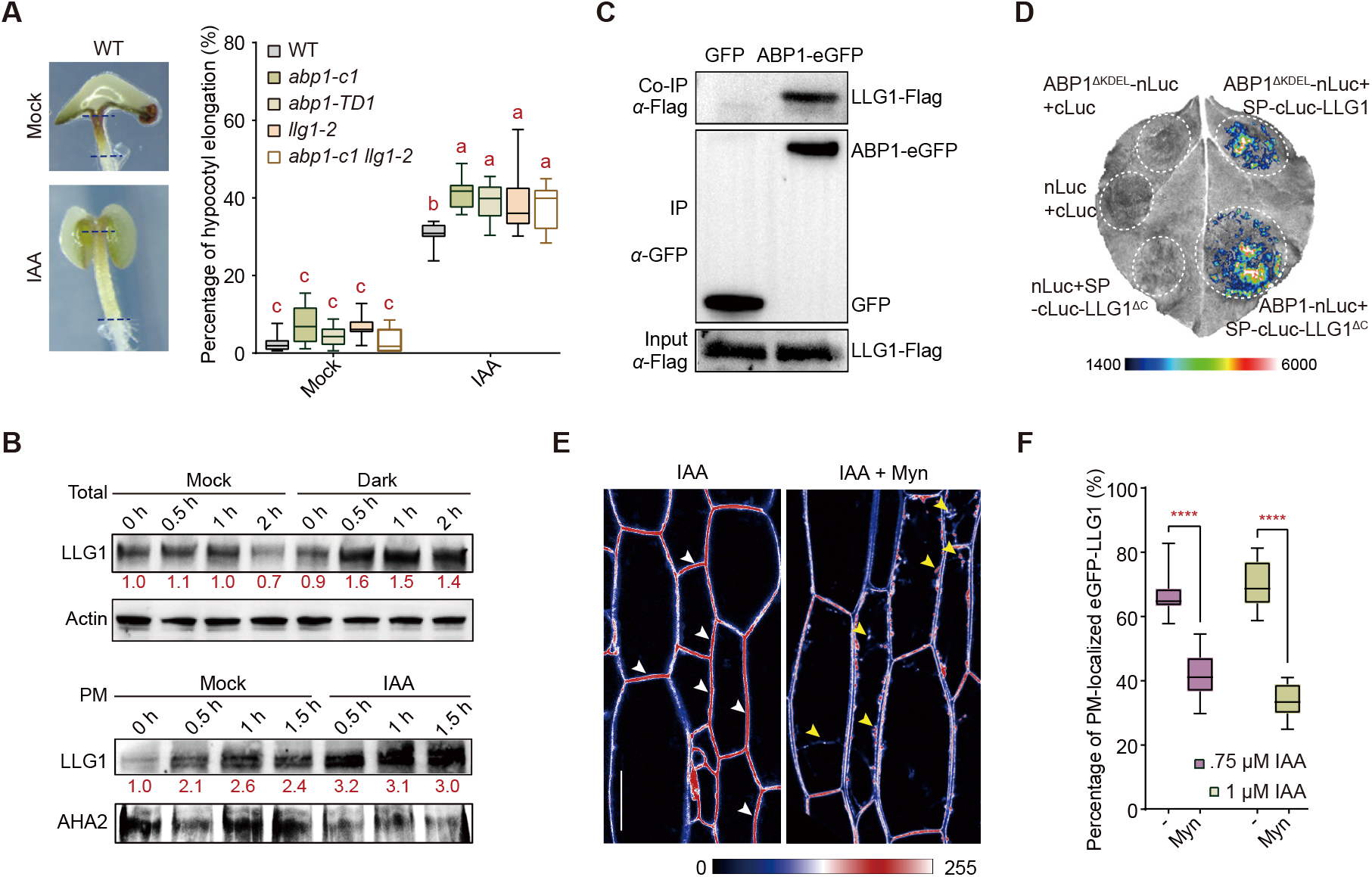
Dark-induced LLG1 expression facilitates ABP1 trafficking through protein-protein interaction. (A) Schematic illustration of the lanolin paste application and representative phenotypes of WT seedlings with or without 1 mM IAA treatment applied to the cotyledons. Quantification of hypocotyl elongation in 4-day-old light-grown WT, *abp1-c1, abp1-TD1, llg1-2* and *abp1-c1 llg1-2* seedlings with or without IAA treatment. *n* ≥ 9 hypocotyls. (B) Western blot analysis of eGFP-LLG1 protein levels under darkness and IAA treatment. Total protein levels of eGFP-LLG1 in 4-day-old seedlings subjected to darkness for the indicated periods are shown (top). Plasma membrane (PM) fractions were isolated and analyzed for eGFP-LLG1 protein levels after IAA treatment for the indicated periods (bottom), with α-AHA2 used as a PM marker. Numbers in red indicate the relative protein amount of eGFP-LLG1 at each time point compared with the non-treated sample. (C) Co-IP of ABP1-eGFP and LLG1-Flag from the *pABP1:ABP1-eGFP pLLG1:sp-mCherry-LLG1-Flag* transgenic plants using anti-Flag antibody. ABP1-eGFP was detected by anti-GFP antibody. (D) Luciferase complementation assay showing the interaction between ABP1^ΔKDEL^ and LLG1 or ABP1 and LLG1^ΔC^. (E) Confocal microscopy imaging of hypocotyl epidermal cells in 4-day-old seedlings treated with IAA alone or Myn co-treatment (pseudo-colored with UnionJack LUT). Scale bar, 20 µm. (F) The percentage of PM-localized eGFP-LLG1 in hypocotyl epidermal cells of 4-day-old seedlings treated with IAA alone or Myn co-treatment. In (A, F), data are presented as mean ± SD. Significant differences were determined using two-way ANOVA (A). Different lowercase letters indicate significant differences at *P* < 0.01. Significant differences were determined using Student’s *t*-test (F). \*\*\*\**P* < 0.0001.

To further investigate the auxin–*LLG1* relationship, we performed an *LLG1* promoter analysis using PlantCARE and identified one auxin-responsive TGA element (fig. S5E). Accordingly, transcription of *LLG1* was induced after 30 min of IAA treatment (fig. S5F). Similarly, LLG1 transcription increased markedly within 2–4 h of the dark treatment (fig. S5G). Consequently, the LLG1 protein levels were markedly increased by auxin and darkness from 0.5 h onward, with high enrichment in the PM fraction (Fig. 2B). We inferred that dark-induced auxin promoted LLG1 accumulation by upregulating its transcription.

Based on these phenotypic and gene expression analyses, we investigated potential interactions between LLG1 and ABP1. Using AlphaFold3, we identified a potential strong interaction interface between ABP1^Δ^^SP^ and LLG1^core^. Residues within this predicted binding region showed predicted Local Distance Difference Test (pLDDT) scores exceeding 70, supporting the reliability of the modeled interface (fig. S6A). Strong interactions were detected by co-immunoprecipitation analysis using *pABP1:ABP1-eGFP* and *pLLG1:mCherry-LLG1-Flag* dual transgenic seedlings (Fig. 2C). Moreover, a luciferase complementation assay (split-Luc) using tobacco leaves demonstrated an interaction between ABP1-nLuc and SP-cLuc-LLG1 (fig. S6B). GPI anchoring requires a C-terminal GPI-attachment signal (*32*); therefore, we generated a GPI-anchoring-deficient LLG1 variant (LLG1^Δ^^C^) and found that deletion of this region did not impair its interaction with ABP1^Δ^^KDEL^, which lacks the ER-retention KDEL motif (Fig. 2D and fig. S6C).

GPI anchor remodeling is required for the delivery of GPI-APs to the PM (*32*). Because Myriocin (Myn) blocks this process in yeast (*33*), and plant GPI-APs contain similar inositol phosphoceramide tails (*34*); we tested whether Myn disrupted GPI-AP function in plants. Root hair growth observations showed that Myn treatment effectively inhibited root hair growth and induced various growth-arrested phenotypes in *llg1-2* (fig. S6D). Myn treatment resulted in obvious intracellular retention of LLG1 in growing root hairs, indicating that inhibition of GPI-AP trafficking impaired proper LLG1 localization (fig. S6E). These results suggest that as a GPI-AP, LLG1 may utilize this trafficking pathway to mediate the targeted localization of specific PM proteins. We found that co-treatment with Myn and IAA significantly attenuated LLG1 localization to the PM (Fig. 2E–F and fig. S6F). In contrast, co-treatment with Myn and IAA did not change the overall LLG1 abundance (fig. S6G). Therefore, we established a Myn treatment system and confirmed trafficking of LLG1 into the PM through a GPI-AP-specific pathway.

### LLG1–ABP1 interaction is auxin dose- and pH-dependent, favoring complex formation under ER-like conditions

To determine whether the distinct pH environments of the ER (∼pH 7.5) and apoplast (∼pH 5.5) affected the ABP1□LLG1 interaction, we performed pull-down and microscale thermophoresis (MST) assays across a pH gradient. ABP1^Δ^^SP^ and LLG1^Δ^^SP^ interacted strongly under ER-like conditions, which progressively weakened as pH decreased (Fig. 3A). Consistently, MST analysis revealed a pronounced pH dependence, with the strongest binding at pH 7.5 *(K*_d_ = 0.044±0.014 μM), substantially weakening at pH 6.5 *(K*_d_ = 4.5±1.21 μM) and being undetectable at pH 5.5 (Fig. 3B and fig. S7A). These results indicate preferential stabilization of the ABP1**–**LLG1 interaction in a neutral ER environment and destabilization in the acidic apoplast.

**Figure 3.**
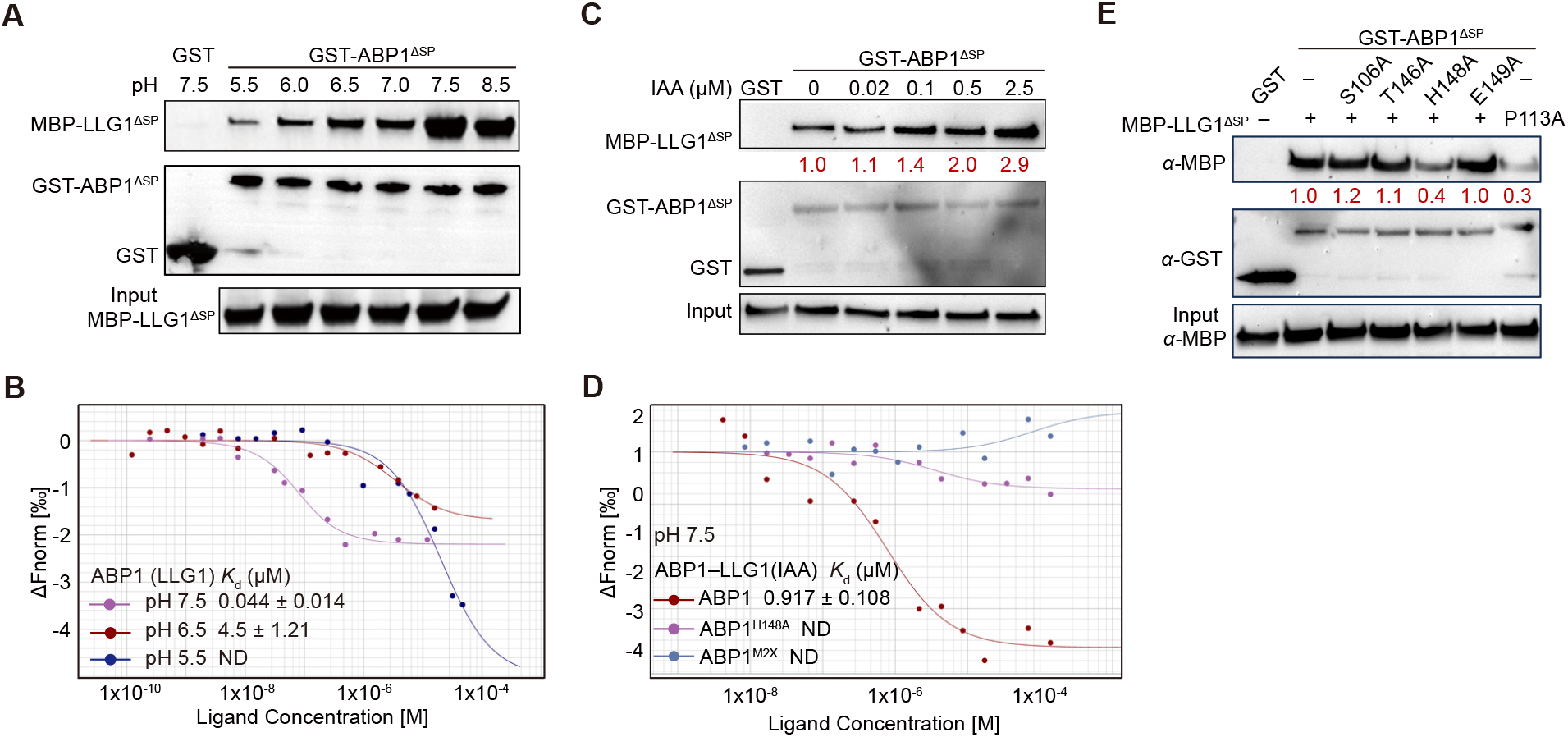
LLG1–ABP1 interaction is auxin dose- and pH-dependent. (A) Pull-down assay showing ABP1^ΔSP^–LLG1^ΔSP^ interaction under pH 5.5–pH 8.5. (B) ABP1–LLG1 binding was measured by microscale thermophoresis (MST) under different pH conditions. Data points indicate the difference in normalized fluorescence (%) generated by ligand-free or LLG1-bound fluorescently labeled ABP1, and the curves show the calculated binding fits. Data are representatives of three independent experiments. ABP1 and LLG1 proteins were expressed and purified from *E. coli* using a prokaryotic expression system. Data are presented as mean ± SD *(n* = 3). ND = Not Detected: No measurable binding signal observed. *K*_d_, dissociation constant. (C) Pull-down assay showing ABP1^ΔSP^–LLG1^ΔSP^ interaction after IAA treatment under pH 7.5. (D) Auxin (IAA) binding to ABP1-LLG1, ABP1^M2X^-LLG1 and ABP1^H148A^-LLG1 was measured by microscale thermophoresis (MST). Datapoints indicate the difference in normalized fluorescence (%) generated by no-liganded or liganded fluorescently labeled LLG1, and the curves show calculated fits. Data are representatives of three independent experiments. ABP1 and LLG1 proteins were expressed and purified from *E. coli* using a prokaryotic expression system. IAA dissolved in ethanol in binding buffer were used for auxin-binding. Data are presented as mean ± SD *(n* ≥ 3). ND = Not Detected: No measurable binding signal observed. *K*_d_, dissociation constant. (E) Pull-down assay demonstrating interactions between: GST-ABP1^Mut^ (S106A, T146A, H148A, E149A) and MBP-LLG1^ΔSP^ (Lane 3–6), GST-ABP1^ΔSP^ and MBP-LLG1^P113A^ (Lane 7) under pH 7.5.

Next, we examined whether auxins modulated this interaction. Using an IAA concentration gradient (0.02–2.5 μM), auxin enhanced the ABP1–LLG1 interaction in a dose-dependent manner at pH 7.5, but not at pH 5.5 (Fig. 3C and fig. S7B–C). MST measurements showed that treatment with 0.5 μM IAA induced a shift in the binding curve of the ABP1–LLG1 interaction at pH 7.5 (fig. S7D). Consistently, MST analysis revealed that IAA could bind to the ABP1–LLG1 complex with a *K*_d_ of 0.917 ± 0.108 μM (Fig. 3D). To determine whether auxin was recognized directly by ABP1, we quantified ABP1–IAA binding and found that ABP1 bound IAA at pH 5.5 with a *K*_d_ of 7.06±2.84 μM, whereas binding at pH 7.5 was below the detection limit (fig. S7E). Furthermore, IAA failed to enhance the interaction between LLG1^Δ^^SP^ and the auxin-binding mutant ABP1^M2XΔ^^SP^ (H59A/H61A) (*10*) at either pH 7.5 or pH 5.5 (fig. S7B–C). MST did not consistently detect auxin-dependent interactions involving ABP1^M2X^ (fig. S7F). These results support a model wherein auxin binding to ABP1 promoted the formation of the ABP1–LLG1 complex.

To identify the residues critical for ABP1–LLG1 interaction, we selected four ABP1 residues (S106, T146, H148, and E149) located at the predicted interface for mutational analysis. Among them, substitution of H148 with alanine markedly reduced the interaction between ABP1 and LLG1^Δ^^SP^. Consistent with the structural model, mutation in the spatially adjacent residue P113 in LLG1 substantially weakened its interaction with ABP1^Δ^^SP^ (Fig. 3E). MST analysis further showed that the IAA-enhanced ABP1–LLG1 interaction was undetectable when H148 was substituted with alanine at pH 7.5 (Fig. 3D). These results indicate that ABP1 H148 and LLG1 P113 are key determinants of the ABP1–LLG1 interface and that the formation of this complex is required for auxin-promoted ABP1 association with LLG1. Phylogenetic analysis revealed high conservation of the ABP1 H148 residue and the ER-retention KDEL motif across vascular plants, and absence in the basal angiosperm *Amborella trichopoda* and non-vascular mosses (fig. S8A). Similarly, the interacting residue LLG1 P113 displayed a nearly identical conservation pattern, retained in vascular plants but not in moss lineages (fig. S8B). This parallel conservation suggests evolution of the receptor trafficking module as an integrated unit in vascular plants.

### LLG1 chaperones auxin-induced ABP1 transport through a GPI-AP trafficking pathway

Biochemical assays demonstrated the function of auxins in ABP1 translocation. We then generated *pABP1:GFP-ABP1*^*M2X*^ and *pABP1:ABP1*^*M2X-GFP*^ plants and found that neither could rescue the defective dark-induced hypocotyl elongation phenotype of the *abp1-c1* mutant (Fig. 4A). Further microscopy analyses revealed that although 1 μM IAA induced the partial translocation of ABP1^M2X^-eGFP to the PM, the extent of this response was markedly weaker than that of wild-type ABP1 (fig. S9A). Thus, the binding of IAA to ABP1 is necessary for the outward secretion of ABP1.

**Figure 4.**
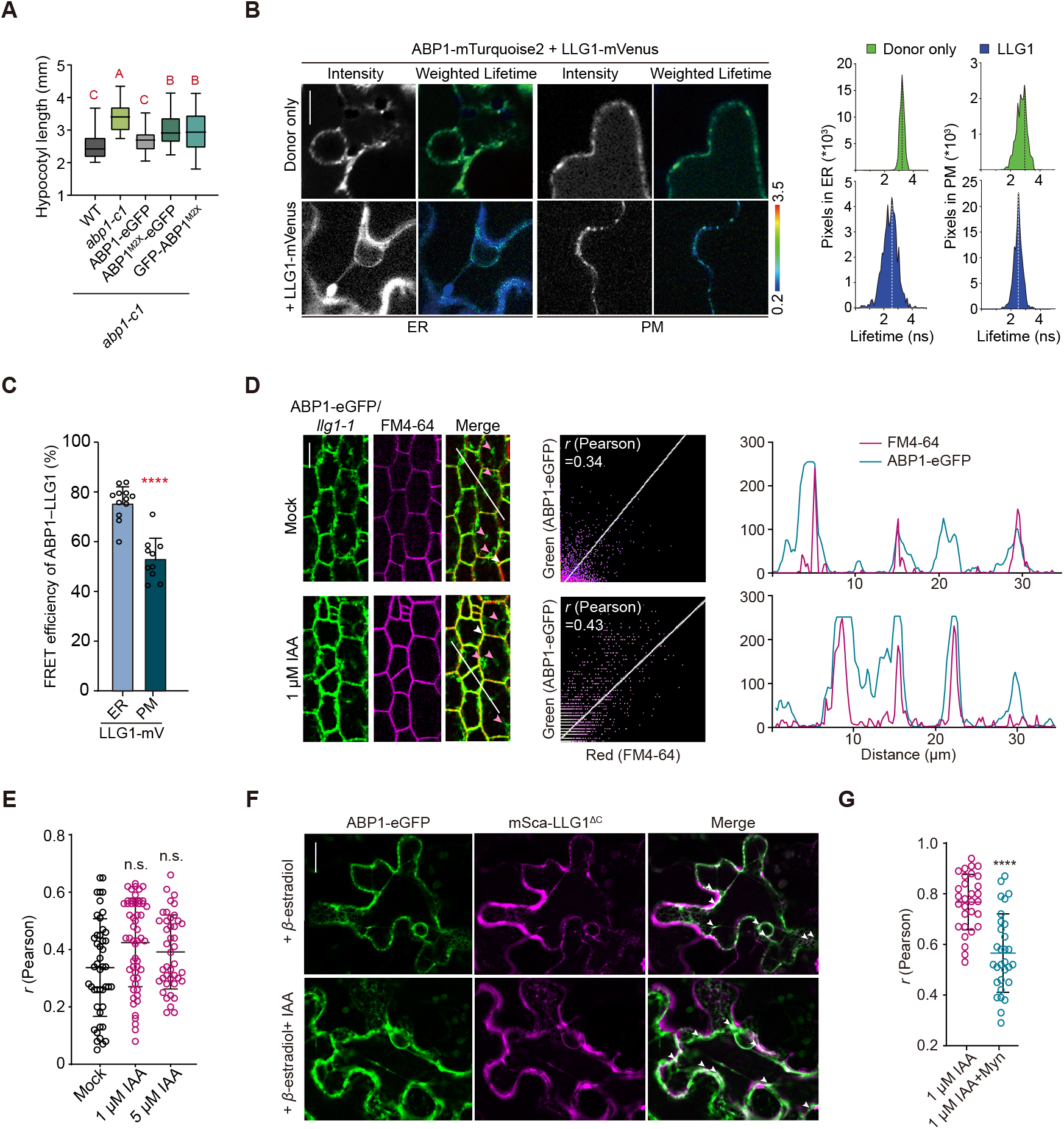
LLG1 facilitates auxin-induced ABP1 trafficking from ER to PM. (A) Quantification of hypocotyl length in 4-day-old WT, *abp1-c1, pABP1:ABP1-eGFP/abp1-c1, pABP1:GFP-ABP1*^*M2X*^*/abp1-c1* and *pABP1:ABP1*^*M2X*^*-eGFP/abp1-c1* transgenic plants subjected to darkness for 36 h. *n* ≥ 45 hypocotyls. (B) Representative FRET-FLIM images (left), lifetime histograms (right) of the ABP1–LLG1 interaction at ER and PM. Intensity-weighted lifetime images use rainbow LUT pseudo-colouring. Histogram dashed lines mark peak lifetime values. Scale bar, 10 µm. (C) Quantification of FRET efficiency of the ABP1–LLG1 interaction at ER and PM. *n* ≥ 10 cells. (D) Confocal microscopy imaging (left), scatterplot analysis with Pearson’s colocalization coefficient (middle) and fluorescence plot profiles (right) of ABP1-eGFP in *llg1-1* background in hypocotyl epidermal cells of 4-day-old seedlings with/without IAA treatment for 1 h. FM4-64 staining indicates the PM, white arrowheads indicate the colocalized signal; pink arrowheads indicate the non-colocalized signal. White lines are used for fluorescence intensity measurement. Scale bar, 10 µm. (E) Pearson’s coefficient showing the colocalization of FM4-64 and ABP1-eGFP in *llg1-1* background after IAA treatment for 1 h. *n* ≥ 42 cells from 15 hypocotyls. (F) Fluorescence imaging of ABP1-eGFP with mSca-LLG1^ΔC^ co-expression in *N. benthamiana* leaves. *pABP1:ABP1-eGFP* and *pER8:mSca-LLG1*^*ΔC*^ in *N. benthamiana* leaves were treated with *β*-estradiol for 24 h and 1 μM IAA treatment for 35 min. White arrowheads indicate the regions of colocalization. Scale bar, 20 µm. (G) Pearson’s coefficient of ABP1-eGFP and FM4-64 colocalization treated with IAA alone or Myn co-treatment. *n* = 30 cells. *(r)*, Pearson’s colocalization coefficient. In (A, C, E, G), data are presented as mean ± SD. Significant differences were determined using one-way ANOVA (A). Different uppercase letters indicate significant differences at *P* < 0.01. Significant differences were determined using Student’s *t*-test (C, E, G). n.s., not significant. \*\*\*\**P* < 0.0001.

To further investigate LLG1-mediated ABP1 trafficking, we performed FRET-FLIM analysis of tobacco leaves expressing ABP1-mTurquoise2 and LLG1-mVenus. Co-expression of LLG1-mVenus markedly reduced the fluorescence lifetime of ABP1-mTurquoise2 in both the ER and PM, with notably higher FRET efficiency in the ER (∼60– 80%) than in the PM (∼40–60%), consistent with their binding affinities (Fig. 4B–C). Notably, in the *llg1-1* mutant, auxin treatment failed to promote ABP1-eGFP translocation to the PM (Fig. 4D–E and S9B). In contrast, using agroinfiltration-mediated expression of *pABP1:ABP1-eGFP* and *sp-mScarlet-LLG1 (XVE)*, IAA treatment induced clear intracellular relocation of the ABP1-eGFP protein. Subsequent β-estradiol induction of *LLG1* promoted ABP1-eGFP transport to the PM, where it colocalized with mScarlet-LLG1 (PM marked by AHA2-eGFP). However, ABP1^H148A^ remained in the ER and failed to reach the PM (fig. S9C–F). These results confirm the role of LLG1 in ABP1 secretion.

Our further analysis of the trafficking mechanism of ABP1–LLG1 revealed that, in contrast to full-length LLG1, the GPI-anchor-deficient LLG1^Δ^^C^ failed to trigger auxin-dependent ABP1-eGFP secretion from the ER to the PM (Fig. 4F). Moreover, co-treatment with Myn suppressed the effect of IAA to promote ABP1-eGFP secretion to the PM, resulting punctate structures reminiscent of the puncta of misfolded GPI-APs resulting from chemically induced ER stress (*35*) (Fig. 4G and fig. S9G–H). We inferred that LLG1 relied on its GPI anchor to facilitate the transport of ABP1. Based on these observations, auxin-induced PM accumulation of ABP1 required approximately 0.5 h. This timescale aligns with the phenotypic manifestations in *abp1* mutants, in which significant differences in hypocotyl elongation rates appear after 2.5 h of dark treatment. We consider these kinetics to be reasonable, given that *de novo* LLG1 synthesis and subsequent GPI-AP-dependent trafficking both play essential roles in ABP1 secretion.

### ABP1 regulates AHA-mediated apoplast acidification in dark-induced hypocotyl elongation

Given the functional association of ABP1 with TMKs and downstream proton ATPases (*9*), we investigated knockout mutants and complementation lines for TMK1/4 and AHA1/2. *tmk1-1* and *tmk4-1* single and double mutants had shorter hypocotyls, which were partially rescued by transgenes encoding either of these two genes (fig. S10A). Nevertheless, both *aha1-7* and *aha2-4* mutants showed defects in hypocotyl elongation, which could be rescued by AHA2 complementation, whereas *ost2-2D*, a constitutively active AHA1 mutant, exhibited a longer hypocotyl than the WT (fig. S10B). Moreover, *aha2-4* and *ost2-2D* exhibited similar responses to auxin and dark treatments (fig. S10C). Further genetic analysis of the *abp1-c1 aha2-4* double mutant revealed a shorter hypocotyl phenotype similar to *aha2-4* (Fig. 5A and fig. S10D). These findings link ABP1 to downstream AHA signaling in dark-promoted hypocotyl elongation. The ABP1/ABL3–TMK1 cell-surface pathway controls PIN2-mediated auxin fluxes required for root gravitropism (*36*). The PIN family protein inhibitor NPA diminished the difference between *abp1* and WT plants, demonstrating that PIN-dependent auxin transport is required for dark-induced hypocotyl elongation (fig. S10E). Therefore, the phenotype of the *tmk1/4* mutant may disturb PIN-mediated auxin efflux and other unknown pathways.

**Figure 5.**
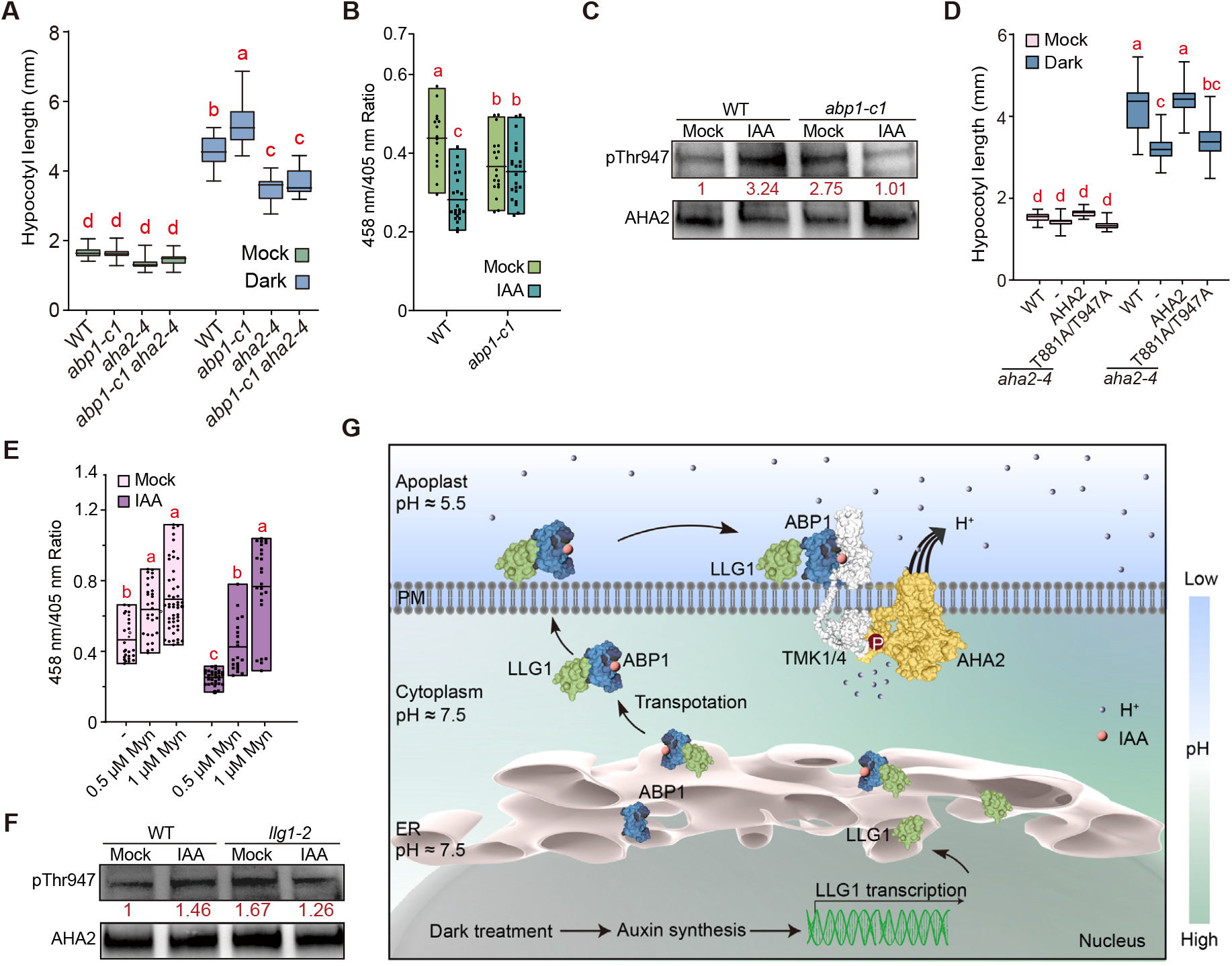
ABP1 regulates the AHAs-mediated dark-induced apoplast acidification and hypocotyl elongation. (A) Quantification of hypocotyl length in WT, *abp1-c1, aha2-4* and *abp1-c1 aha2-4* seedlings subjected to darkness for 30 h. *n* ≥ 23 seedlings. (B) Relative quantification of HPTS-stained apoplastic pH in hypocotyl cortex cells in 4-day-old WT and *abp1-c1* seedlings after dark treatment for 18 h following 1 µM IAA treatment for 15 min. *n* ≥ 15 cells. (C) Western blot detection of phosphorylated AHA2 in WT and *abp1-c1* seedlings (anti-pThr947). AHA2 protein levels were determined using anti-AHA2 antibodies. Seedlings were subjected to darkness for 12 h followed by 1 µM IAA treatment for 10 min. (D) Quantification of hypocotyl length in *aha2-4, pAHA2:AHA2/aha2-4* and *pAHA2:AHA2*^*T881A/T947A*^*/aha2-4* seedlings subjected to darkness for 30 h. *n* ≥ 20 seedlings. (E) Relative quantification of HPTS-stained pH in the apoplast of hypocotyl cortical cells in 3-day-old WT seedlings treated with Myn for 1 d followed by 18 h dark treatment and then 1 μM IAA treatment for 15 min. *n* ≥ 22 cells. (F) Western blot detection of phosphorylated AHA2 in WT and *llg1-2* seedlings (anti-pThr947). AHA2 protein levels were determined using anti-AHA2 antibodies. Seedlings were subjected to darkness for 12 h followed by 1 µM IAA treatment for 10 min. (G) Model of dark/auxin-induced and LLG1-facilitated ABP1 trafficking in hypocotyl elongation. Dark treatment induces auxin biosynthesis and subsequently stimulates *LLG1* expression. Auxin enhances LLG1 interaction with ABP1 in the ER and assists its trafficking through GPI-APs unique pathway. Upon reaching the PM, ABP1 dissociates from LLG1 under apoplastic acidic conditions and in turn initates the TMKs–AHAs-mediated signaling. In (A, B, D, E), data are presented as mean ± SD. Significant differences were determined using two-way ANOVA. Different lowercase letters indicate significant differences at *P* < 0.05.

To further explore AHA1/2-involved downstream signaling, we conducted a pH indicator HPTS staining assay. Apoplast pH induced by darkness in the WT hypocotyl epidermis and cortex was significantly reduced and was more acidic in *abp1-c1* (Fig. 5B and fig. S11A). In line with this, auxin treatment followed by darkness induced rapid and pronounced acidification in the WT but not the *abp1-c1* mutant (fig. S11B–D). Nevertheless, no obvious differences were detected under light conditions (fig. S11E–F). Consistent with the well-established function of AHA2 Thr947 phosphorylation in pump activation (*37*), auxin/dark treatment failed to induce this phosphorylation level in *abp1-c1* mutant compared to the WT (Fig. 5C). Moreover, the mutation of the two established AHA2 activation residues (T881 and T947) (*3, 38*) abolished the ability of pAHA2/T947A to rescue hypocotyl elongation defects in *aha2-4* mutant (Fig. 5D).

In addition, we applied Myn pretreatment before darkness and auxin treatment to investigate LLG1 participation. Notably, Myn application significantly disturbed the auxin-induced apoplast acidification in the hypocotyls (Fig. 5E and fig. S11G). Furthermore, upon auxin treatment in the dark, the auxin-induced Thr947 phosphorylation response was absent in *llg1-2*, mimicking the phenomenon observed in the *abp1* mutant (Fig. 5F). These results establish a causal link between LLG1-facilitated ABP1 transport and AHA2-mediated apoplast acidification.

## Discussion

Our findings establish a darkness induction system that stimulate auxin synthesis and upregulates LLG1 expression in hypocotyl. Auxin promotes the interaction between LLG1 and ABP1 in a dose-dependent manner and facilitates vesicular trafficking via the GPI-AP-specific pathway. Upon reaching the apoplast, ABP1 dissociates from LLG1 owing to the acidic apoplastic pH and subsequently coordinates with the AHA-mediated pathway for the regulation of hypocotyl elongation (Fig. 5G).

In contrast to previous observations that ABP1 exhibits little detectable auxin binding under ER-like conditions (*20, 39*), we identify LLG1 as a critical cofactor that facilitated ABP1-mediated auxin perception in the ER. Auxin promotes the formation of the ABP1–LLG1 complex, thereby coupling hormone sensing with receptor mobilization and subsequent cell surface signaling. These findings provide a mechanistic explanation for how auxin overcomes the ER retention of ABP1 and establish a framework in which ABP1 transitions from an intracellular sensing state to a signaling-competent state in the apoplast. Moreover, the pH-sensitivity of the ABP1–LLG1 interaction likely arises from the reversible protonation of ABP1 His148, which may remodel the local electrostatic interface, thereby promoting complex dissociation in the acidic apoplast (*40, 41*). The coordinated conservation of H148, the KDEL motif, and LLG1 P113 across vascular plants suggests that coupling intracellular auxin perception with cell surface signaling is an evolutionarily conserved feature of rapid auxin responses.

Our findings identify ABP1 as an intracellular auxin receptor whose signaling activity depends on regulated release from ER retention. In eukaryotes, the C-terminal KDEL motif mediates retrieval from the Golgi apparatus to the ER, thereby limiting secretion (*42, 43*). Herein, we showed that the auxin-induced association with LLG1 facilitated ABP1’s escape from this retention and reached the apoplast. We propose that the abundance of LLG1-bound ABP1 exceeds the capacity for KDEL receptor-mediated retrieval, thereby enabling ABP1 delivery to the apoplast for rapid signaling. Notably, this mode of transport differs from the conventional vesicular trafficking used by most membrane and secretory proteins, as GPI-APs are sorted through sphingolipid-dependent membrane microdomains, a conserved pathway that enhances trafficking efficiency across eukaryotes (*44-46*). Moreover, as a GPI-AP, LLG1 may facilitate the lateral organization of ABP1 on the PM (*47, 48*), thereby promoting its interaction with TMK receptor kinases and subsequent AHA-mediated auxin signaling.

Our findings establish a conceptual framework in which GPI-AP-mediated trafficking links intracellular receptor retention to signal activation, potentially representing a broadly conserved mechanism in eukaryotes.

## Acknowledgments

We express our sincere gratitude to Dr. Lin Li and the Large-Scale Instrument Sharing Platform of the School of Life Sciences at Fudan University for support with the hypocotyl growth kinetic analysis, and Dr. Peng Zhang and Dr. Xue Zhang for assistance with the structural prediction analysis. We thank Dr. Jiří Friml for providing the seeds of *pABP1:GFP-ABP1-M2X*/*abp1-c1*, Dr. Lin Xu for *abp1-c1* and *abp1-TD1*, Dr. Jun Liu for *ost2-2D*, Dr. Meng-xiang Sun for RFP-HDEL and Dr. Peng Zhao for mScarlet-K32 vector. We also acknowledge the Institute of Genetics and Developmental Biology, CAS for the IAA quantification assay, the Materials Characterization Center, and the Instruments Sharing Platform of the School of Life Sciences, East China Normal University.

## Funding

National Natural Science Foundation of China grant 32425008, 32230009 (CL), 32500276 (BYL). Science and Technology Commission of Shanghai Municipality grant 24N12800100 (CL).

## Author contributions

Conceptualization: CL

Methodology: JW, CL, MMZ, BYL, YZH, YQY Investigation: JW, JPY, MMZ, HQF

Project administration: CL, JW, BYL, HQF Supervision: CL

Writing – original draft: CL, JW, JPY Writing – review & editing: CL, TDX

## Competing interests

Authors declare that they have no competing interests.

## Supplementary Materials

Materials and Methods

Figs. S1 to S11

Tables S1

References (*49–63*)

## Notes

### Competing Interest Statement

The authors have declared no competing interest.

### Summary of Updates

Figure 3 revised; author updated; Supplemental files updated. Plate sorting has been revised.

