## Supplemental S1-11 for "Receptor trafficking couples intracellular auxin perception to rapid signaling"

### Materials and Methods

#### Plant materials and growth conditions

*Arabidopsis* ecotype Col-0 was used as wild type (WT) in this study. The *abp1-c1*, *abp1-TD1 (49)*, *llg1-2*, *aha1-7*, *aha2-4*, *ost2-2D(50)* and *pYUC9:GUS (51)* have been described previously. Seeds were surface-sterilized with 75% (v/v) ethanol for 5 min, followed by five times washes with sterilized distilled water. The sterilized seeds were then germinated on plates containing half-strength Murashige and Skoog (1/2 MS) medium supplemented with 1% (w/v) sucrose and solidified with 0.8% (w/v) agar, adjusted to pH 5.7. *Arabidopsis* seedlings were cultivated in Percival growth chambers under controlled environmental conditions (hereafter, standard growth conditions): 16 h light/8 h dark photoperiod with 22°C light/20°C dark temperatures and 60% relative humidity. The light regimes were implemented under regular white light at an intensity of 100 $\mu mol \cdot m^{-2} \cdot s^{-1}$.

#### Dark treatment and hypocotyl growth assay

The kinetics of hypocotyl growth were monitored using the high-throughput imaging platform DynaPlant®. Seeds were sown on 1/2 MS medium solidified with 2% phytagel (Solarbio, P8170). 3-day-old seedlings were pretreated under continuous white light for 24 h for acclimation to fresh medium, and subsequently transferred to complete darkness for 36 h. During dark incubation, images were captured at 15 min intervals (spatial resolution of 1.2 µm per pixel) and hypocotyl growth lengths in the time-series images were quantified by DynaPlant Analysis software (*22*). Data are presented as mean ± SEM. Means were calculated from hypocotyls of more than 28 seedlings. Experiments were performed with at least three biological replicates, with each replicate derived from distinct biological samples to minimize batch effects.

Hypocotyl imaging was conducted using a stereomicroscope (Leica M205 FCA) and a fluorescence microscope (Leica DM2500) equipped with a 20× water-immersion objective. Hypocotyl length was measured in ImageJ from the axillary bud to the hypocotyl-root junction. For cellular analyses, individual cell areas were quantified by tracing cell boundaries using the polygon selection tool in ImageJ. 4-Phenoxyphenyl-boronic acid (*52*) (PPBo, MedChemExpress, HY-76584) or N-1-Naphthylphthalamic acid (*53*) (NPA, MedChemExpress, HY-116425) was added to solid 1/2 MS medium at specified concentrations for dark treatment. 4-day-old seedlings grown under standard conditions were transferred to the respective plates, which were then wrapped in two layers of aluminum foil to ensure darkness and placed vertically in Percival growth chambers. Hypocotyl imaging and length measurement were performed as described above.

#### Auxin treatment and hypocotyl length assay

Seedlings were treated with liquid 1/2 MS medium (1% sucrose, pH 5.7), supplemented with indicated concentrations of indole-3-acetic acid (IAA, Sigma-Aldrich, I2886). All treatments were performed in 1.5 mL Eppendorf tubes under standard growth conditions with continuous white light (~100 $\mu mol \cdot m^{-2} \cdot s^{-1}$). For IAA treatment of cotyledons, lanolin (Aladdin, L116309) was used as the medium, as outlined in established protocols (*31*). Briefly, lanolin was melted at 65°C, mixed with 1mM IAA (at the desired concentrations) or DMSO, and homogenized thoroughly. Seedlings were then vertically positioned (adaxial-side down) in the lanolin-IAA mixture to ensure uniform cotyledon coating before solidification. The treated seedlings were then transferred to 1/2 MS solid medium and cultured vertically under standard growth conditions. The means were calculated from at least 9 hypocotyls. Experiments were performed with at least three biological replicates, with each replicate derived from distinct biological samples to minimize batch effects.

#### Free IAA level measurement

Seedlings were grown for 4 days under standard conditions and then transferred to darkness for 0, 12 or 24 h. For the quantification of phytohormone IAA, ~200 mg of cotyledon-hypocotyl tissues mixtures were harvested, spiked with 3 ng [$^{13}C_6$] IAA internal standard and processed through extraction, purification, and methylation prior to analysis by selected-ion-monitoring GC-MS following established protocols (*54*). The measurement was performed by the Institute of Genetics and Developmental Biology, Chinese Academy of Sciences.

#### Histochemical GUS staining assay

For GUS staining assay, seedlings were incubated in buffer containing 100 mM sodium phosphate buffer (pH 7.0), 10 mM EDTA, 0.1% (v/v) Triton X-100, 0.5 mM potassium ferricyanide, 0.5 mM potassium ferrocyanide and 1 mM X-Gluc at 37°C in darkness for 12 h, followed by incubation in 70% ethanol for 5 h with the ethanol replaced every hour. The samples were photographed under a stereoscope (Leica M205 FCA).

#### Plasmid construction and transgenic plants generation

Primers used in this study can be found in table S1. Full-length ABP1 and LLG1 cDNAs were PCR-amplified from WT for subsequent cloning. For plant transformation, the genes were cloned into *pCAMBIA1300* to generate *pLLG1:sp-eGFP-LLG1* and *pABP1:ABP1-eGFP* constructs, which were subsequently introduced into *llg1-2* mutant and WT plants respectively using *Agrobacterium tumefaciens*-mediated floral dip transformation (*55*). For protein interaction studies, the cDNAs were first cloned into *pDONR221* vectors (*P3P2-ABP1* and *P1P4-LLG1*) using BP recombination, then recombined into the *pFRETtv-2in1-CC7* vector (Addgene 105116) to create the FRET reporter construct *pFRET-ABP1-mTurquoise2–LLG1-mVenus*. Additionally, ABP1 was subcloned into *pGEX-6P-1* for prokaryotic expression in *E. coli* (Rosetta, BL21).

For *SP-cLuc-LLG1*, sequences encoding the signal peptide of LLG1 (residue 1–23), nLuc, and the deleted signal peptide of LLG1 (residues 24–168) were amplified by PCR and cloned into *pCAMBIA1300* driven by the 35S promoter using a ClonExpress II One Step Cloning Kit. For *SP-cLuc-LLG1*$^{\Delta GPI}$, sequences encoding the signal peptide of LLG1 (residue 1–23), nLuc, and the deleted cytoplasmic domain of LLG1 (residues 24–147) were amplified. For *ABP1-nLuc*, sequences encoding the ABP1 (residues 1–198) was fused with cLuc. For *ABP1*$^{\Delta KDEL}$, sequences encoding the ABP1 (residues 1–194) was fused with cLuc. Primers used in plasmid construction are listed in table S1.

For *sp-mScarlet-LLG1 (XVE),* sequences encoding the signal peptide of LLG1 (residue 1–23), mScarlet, and the deleted signal peptide of LLG1 (residues 24–168) were amplified by PCR and cloned into *pER8* vector of the *XVE* inducible system (*56*) using a ClonExpress II One Step Cloning Kit. In *Nicotiana benthamiana* leaves, estradiol-activated *XVE* induces expression of the *mScarlet* reporter gene driven by a target promoter consisting of eight copies of the LexA operator fused upstream of the −46 35S minimal promoter.

For *pAHA2:AHA2/aha2-4*, full-length *AHA2* cDNA was PCR-amplified from WT seedlings for subsequent cloning. To generate the native promoter-driven complementation construct, the *AHA2* genomic fragment containing the promoter region and coding sequence in *pCAMBIA1300* produced *pAHA2:AHA2.* For the phospho-deficient mutant construct *pAHA2:AHA2*$^{T881A/T947A}$*/ aha2-4*, site-directed mutagenesis was performed to substitute Thr881 and Thr947 with alanine residues (T881A and T947A). The mutated AHA2 fragment was then cloned into *pCAMBIA1300* under the control of the native AHA2 promoter to generate *pAHA2:AHA2*$^{T881A/T947A}$. The resulting constructs were introduced into the *aha2-4* mutant background by floral dipping, and transgenic lines were selected on hygromycin-containing medium for further analysis.

**Plant protein extraction and western blot**

For the detection of changes in protein levels, plant tissue samples from specific periods and treatments were harvested and ground in liquid nitrogen. Total proteins were extracted using Tris-HCl buffer (pH 6.8) through sequential centrifugation (15,000 × g, 30 min; then 15,000 × g, 10 min) at 4°C. For MG132 (MedChemExpress, HY-13259) treatment, it was diluted with 1/2 MS liquid medium to 30 μM and applied 30 min prior to IAA treatment. Protein samples were quantified by BCA assay, mixed with loading buffer, and heat-denatured at 99°C for 10 min.

PM proteins were extracted using a commercial kit (Solarbio, China, EX2080). Protease inhibitor cocktail (2 μL) was added to 500 μL of cold Extraction Buffer A, mixed thoroughly, and kept on ice. Plant tissue (100–200 mg) was finely chopped and homogenized in 500 μL of Extraction Buffer A at 2–8°C. The homogenate was transferred to a pre-cooled centrifuge tube, shaken at 2–8°C for 30–60 min, and then centrifuged at 12,000 × g for 5 min at 2–8°C. The supernatant was collected, mixed with 5 μL of Extraction Buffer B, incubated at 37°C for 10 min, and centrifuged at 1,000 × g for 3 min at 37°C. After removing the upper layer, the remaining 30–50 μL lower phase was dissolved in 50–150 μL ice-cold Membrane Protein Solubilization Buffer C to obtain the membrane protein fraction.

ER proteins were extracted using a commercial kit (Solarbio, China, EX2061). Briefly, protease inhibitor cocktail was added to Extraction Buffer C (2 μL per 300 μL), mixed, and kept on ice. Approximately 200–500 mg of plant tissue (de-veined and cleaned) was finely chopped, homogenized in 1 mL Extraction Buffer A using a homogenizer or Dounce homogenizer, and filtered through a 100 μm cell strainer. The filtrate was sequentially centrifuged at 200 × g for 3 min, 500 × g for 5 min, 1000 × g for 10 min, and 20,000 × g for 10 min, collecting the supernatant at each step. This was followed by ultracentrifugation at 50,000 × g for 45 min to pellet microsomal membranes (using a benchtop ultracentrifuge, Thermo Sorvall MTX150). The pellet was resuspended in 400 μL of cold Reagent B, centrifuged again at 50,000 × g for 45 min, and the resulting pellet was resuspended in 200–300 μL of Extraction Buffer C. After shaking at 2–8 °C for 30 min, the mixture was centrifuged at 12,000 × g for 10 min at 4 °C. The supernatant containing ER proteins was collected, quantified, aliquoted, and stored at −80 °C for further use.

For Western blot detection, anti-GFP (1:2000 dilution, sc-9996, Santa Cruz, America) detected eGFP-LLG1 or ABP1-eGFP fusion protein expression. Anti-Bip (1:2000 dilution, Agrisera, AS09481, Sweden) served as ER reference proteins. Anti-AHA2 (1:1000 dilution, PY231, Beijing QiWei YiCheng Tech Co., Ltd) served as PM reference proteins. HRP-conjugated Goat Anti-Mouse/Rabbit IgG (1:5000; Sangon Biotech D110087/D110058)

**Promoter element analysis**

Promoter element analysis is carried out using PlantCARE (*57*), a database of plant promoters and their cis-acting regulatory elements: https://bioinformatics. psb.ugent.be/webtools/plantcare/ html/.

**RNA isolation, RT-PCR, and qRT-PCR analysis**

Seedlings at appropriate developmental stages were harvested, and roots were excised to retain only aboveground parts for transcript profiling of *ABP1* and *LLG1/LRE* genes. RNA was extracted using the RNAprep Pure Plant Kit (TIANGEN, China), and first-strand cDNA was synthesized with a HiScript ®II 1st Strand cDNA Synthesis kit (Vazyme, China). qRT-PCR was performed with ChamQ Universal SYBR qPCR Master Mix (Vazyme, China) using the following thermal profile: 95°C for 5 min, 30 cycles of 95°C for 10 s and 60°C for 30 s. Relative transcript levels were calculated using the $2^{-\Delta\Delta CT}$ method, normalized to *ACTIN2*. Primer sequences are listed in table S1.

#### Confocal microscopy imaging

For colocalization analysis in *A. thaliana* and *N. benthamiana*, images were acquired by the Ultra High Resolution Laser Leica STELLARIS 8 confocal microscope with WLL laser, power HyD detector (HyDx, HyD2 and HyD3) and frame scanning mode. The real fluorescence signal was determined using TauGating (time-gated fluorescence lifetime filtering) and photon counting techniques to suppress autofluorescence, and enhance signal-to-noise ratio (*58*). No fluorescence signal was detected in WT seedlings under identical imaging settings and channels. Intracellular ER was visualized with RFP-HDEL as a marker, while PM was stained using the amphiphilic dye FM4-64 (*59*), allowing simultaneous visualization of both compartments. Fluorescence parameters were as follows: GFP (Ex 490 nm, Em 505–560 nm), RFP (Ex 532nm, Em 550–700 nm), FM4-64 (Ex 523nm, Em 550–700 nm), mScarlet (Ex 594, Em 628 nm). All imaging sessions (including mock controls and experimental samples treated with IAA or NAA [Sigma-Aldrich, N0640]) were performed under identical conditions: 20% WLL laser power, 21°C, and 35% humidity. Statistical analysis was based on >3 cells in approximately 3–4 independent hypocotyls. Each cell area included the complete cell edge, and the same calculation parameters and procedures were used for all cells. Experiments were performed with at least three biological replicates, with each replicate derived from distinct biological samples to minimize batch effects. Myriocin (Myn, GLPBIO, GC14278, Montclair, CA, USA) was added to the solid medium for cultivating *A. thaliana* seedlings. 10 µM *β*-estradiol (Sigma-Aldrich E2758) was pressure-infiltrated 24 h post-transformation, with phenotypic analysis conducted 24 h post-induction.

#### Colocalization analysis

Colocalization was quantified in Fiji software using the Coloc 2 plugin (Analyze › Colocalization Analysis › Coloc 2), yielding colocalization scatter plots, Pearson coefficients. For each sample, paired green and red channel images were loaded, and regions of interest (ROIs) were selected for analysis. Pearson's correlation coefficients (*r*), ranging from −1 (perfect negative correlation) to +1 (perfect positive correlation) (*60*), were calculated to quantify the degree of signal colocalization. Pearson's correlation coefficients (*r*) were computed as follows:

$$r = \frac{\sum (R_i - R_{av}) \cdot (G_i - G_{av})}{\sqrt{\sum (R_i - R_{av})^2 \cdot \sum (G_i - G_{av})^2}}$$

$R_i$, $G_i$ represent the intensity values of the i-th pixel in the red channel ($R_i$) and green channel ($G_i$). $R_{av}$, $G_{av}$ represent the mean intensities of the entire dataset for the red channel ($R_{av}$) and green channel ($G_{av}$). To obtain the colocalization peak plots, merged dual-fluorescence images were converted to color channels: Image > color > channel tool > color channel. A straight line was used to mark the region of interest, and the plot profile's live mode was employed to display the fluorescence peak plots of different channels. The data were saved separately and graphed in GraphPad Prism.

#### Protein structure prediction

Protein structure prediction was performed using AlphaFold3 (v.3) (*61*) in multimeric mode with default parameters. Structural models were visualized using PyMOL (v.3.0.3) (Shelton, 2025), and contact plots were generated using custom scripts.

**Luciferase complementation assay**

*A. tumefaciens* strain GV3101 harboring various constructs was cultured in LB liquid medium until reaching an $OD_{600}$ of 1.0. The cultures were then centrifuged at 3,824 × g for 10 min, and the resulting pellets were washed and resuspended in $ddH_2O$ supplemented with 10 mM MES (pH 5.7), 10 mM $MgCl_2$, 0.01% (v/v) Tween-20 and 100 μM acetosyringone to a final $OD_{600}$ of 0.6. Equal volumes of bacterial suspensions containing the nLUC and cLUC constructs were mixed and infiltrated into designated regions of *N. benthamiana* leaves. Plants were kept in darkness for 12 h, then transferred to a 16 h light/8 h dark cycle for an additional 48 h. To assess luciferase activity, infiltrated leaves were treated with 1 mM luciferin solution [0.05% (v/v) Triton X-100] and incubated in darkness for 5 min, followed by imaging using a Tanon-5200 Multi chemiluminescence/fluorescence imaging system (Tanon, China) equipped with a cooled CCD camera.

**Protein expression and purification**

DNA sequences encoding LLG1 (residues 24–168) and ABP1 (residues 34–198) were cloned into the *pMAL-c5x* or *pGEX-6P-1* vectors for expression in *E. coli* BL21 Competent Cells. For maltose-binding protein (MBP)- and glutathione S-transferase (GST)-tagged proteins, cells were incubated at 37°C with shaking at 200 rpm until the $OD_{600}$ reached 0.6. Following induction with 0.5 mM isopropyl-*β*-d-thiogalactopyranoside (IPTG), cells were incubated with shaking for 3 h at 37°C (MBP-LLG1) or 16 h at 25°C (GST-ABP1), and harvested by centrifugation at 3,824 × g for 10 min at 4°C. Cells were lysed by sonication on ice in lysis buffer containing 50 mM Tris-HCl pH 7.5, 100 mM NaCl, 1 mM $Na_2$-EDTA, 5% (v/v) glycerol , and 1 mM PMSF, and centrifuged at 15,000 rpm for 30 min at 4°C. The supernatants were purified using amylose resin (YESEN, China) for MBP fusion protein and glutathione agarose resin (YESEN, China) for GST fusion protein.

**Pull-down assay**

For the GST pull-down assay, purified MBP-LLG1$^{\Delta SP}$ and GST-ABP1$^{\Delta SP}$ (GST-ABP1$^{\Delta SP\ S106A}$, GST-ABP1$^{\Delta SP\ T146A}$, GST-ABP1$^{\Delta SP\ H148A}$) proteins were resuspended in binding buffer (50 mM Tris-HCl pH 5.5/6.0/6.5/7.0/7.5/8.5, 100 mM NaCl, 1 mM $Na_2$-EDTA, 50 μM $ZnCl_2$, 5% (v/v) glycerol, 1 mM PMSF) and incubated with GSTSep Glutathione Agarose Resin (Yeasen, 20507ES10) at 4°C for 2 h with rotation. The resins were then washed three times in pull-down buffer, followed by the addition of SDS-PAGE loading buffer and boiled for 5 min, and used for immunoblot analysis. The protein blot was stained with Ponceau S for loading detection. Anti-MBP antibody (D195305-0100, 1:2,000 dilution, Sangon Biotech, China), anti-GST horseradish peroxidase-conjugated antibody (B-14, 1:1,000 dilution, Santa Cruz Biotechnology, USA), and anti-mouse secondary antibody (ASS1007, 1:5,000 dilution, Abgent, USA) were used for chemiluminescence detection. The signals were acquired using the Tanon-2500R imaging system (China).

**MicroScale Thermophoresis (MST)**

For MST-based protein interaction analysis, LLG1-MBP and GST-ABP1 fusion proteins were expressed in *E. coli* at 25°C for 16 h and subjected to affinity purification. The harvested bacterial cells were resuspended in lysis buffer [1× PBS containing 1 mM DTT, 1 mM EDTA, and 1 mM PMSF], followed by ultrasonication and 0.5% Triton X-100 addition. The lysate was centrifuged at 8,500 rpm for 30 min and applied to a resin column for protein binding. After washing with 10 column volumes of 1× PBS buffer, the proteins were eluted. Specifically, for GST-ABPs purification, the elution buffer consisted of 10 mM reduced glutathione, 50 mM Tris-HCl (pH 7.5), and 1 mM DTT; whereas for LLG1-MBP purification, the elution buffer contained 10 mM maltose, 20 mM Tris-HCl (pH 7.5), 1 mM EDTA, and 1 mM DTT. Protein concentration was determined using the bicinchoninic acid (BCA) assay.

Protein labeling was performed using the RED-NHS Protein Labeling Kit (Nano Temper, MO-L011) following the manufacturer's instructions. Protein samples with a degree of labeling (DOL) between 0.5 and 1.0 fluorophores per protein were used for subsequent MST measurements. Ligand-protein interaction assays were performed in their respective binding buffers at 4°C for 30 min to ensure equilibrium binding. Reactions were carried out in buffers containing 250 mM NaCl, 50 µM $ZnCl_2$, and 0.01% (w/v) Tween-20, with 50 mM citrate buffer at pH 5.5 or HEPES buffer at pH 6.5 or 7.5. Subsequently, 4–6 µl of each reaction mixture was loaded into pre-treated silica capillaries (Monolith™ NT.115 Standard Treated Capillaries, MO-K002). MST analyses were performed on Monolith NT.115 instrument with settings optimized to 20% LED excitation power and 40% MST laser power. All measurements were replicated at least three times using independently prepared protein samples to ensure technical reproducibility. Equilibrium dissociation constants ($K_d$) were calculated by fitting the binding isotherms to the mass action law using NanoTemper MO. Affinity Analysis software (MO.Control 2).

**FRET-FLIM assay**

*N. benthamiana* leaves were agroinfiltrated to co-express ABP1-mTurquoise2 (donor) and LLG1-mVenus (acceptor) constructs. Imaging was performed 48–72 h post-infiltration on epidermal cells exhibiting dual fluorescence signals. Confocal laser scanning microscopy (CLSM) settings were as follows: excitation 458 nm/emission 465 to 505 nm (mTurquoise2), excitation 514 nm/emission 520 to 560 nm (mVenus). Cells of interest were selected using a 40× water-immersion objective. ABP1-mTurquoise2 was used as the donor only. The fluorescence lifetime of the donor (mTurquoise2) was measured using the FLIM (*62*) module (PicoHarp 300 TCSPC system), with the average value obtained from three measurements as the initial fluorescence lifetime of the donor. The efficiency of mTurquoise2–mVenus FRET was analyzed using a STED Leica microscope with the FRET module. The region of interest (ROI) was selected to focus on the specific subcellular regions, and the initial fluorescence lifetime value of the donor was inputted. FRET efficiency was automatically calculated by the system using the following formula:

$$E_{(\mathrm{FRET-FLIM})} = 1 - \frac{\tau(\mathrm{quench})}{\tau} = 1 - \frac{\tau(\mathrm{donor, FRET})}{\tau(\mathrm{donor, no\ FRET})}$$

τ (donor, FRET) represents the donor lifetime in the presence of the acceptor, and τ (donor, no FRET) represents the donor lifetime in the absence of the acceptor (*62*).

**HPTS staining and imaging**

HPTS staining and imaging were conducted according to standard procedures in the field, with adjustments made to suit the specific requirements of this study (*63*). Prior to staining, hypocotyls of 4-day-old light-grown seedlings or seedlings subjected to 18 h of darkness were equilibrated in 1/2 MS growth medium supplemented with 0.2 mM MES for at least 30 min. For auxin treatment, hypocotyls were incubated in 1/2 MS growth medium supplemented with 5 mM HPTS (prepared from a 100 mM aqueous stock solution) and 1 µM IAA for 15 min. For Myn treatment, seedlings were grown on 1/2 MS medium under continuous light for 3 days and then transferred to 1/2 MS solid medium supplemented with different concentrations of Myn for pre-incubation for 24 h. Subsequently, seedlings were subjected to co-treatment with Myn under dark conditions for 18 h, after which hypocotyls were incubated in 1/2 MS growth medium containing 5 mM HPTS and 1 μM IAA for 15 min. After staining, hypocotyls were mounted in the same growth medium on microscopy slides and covered with coverslips for imaging.

Hypocotyl imaging was performed using an Ultra High-Resolution Leica STELLARIS 8 confocal microscope equipped with a 20× water-immersion objective. Fluorescence signals from protonated and deprotonated HPTS forms were detected using excitation at 405 nm with an emission peak at 514 nm, while signals from the deprotonated form were detected using excitation wavelengths of 405 nm and 458 nm respectively, both with emission peaks at 514 nm. The 458/405 ratio was calculated for extracellular pH quantification. Approximately 5–6 independent hypocotyls were analyzed per treatment, with about 2–3 cells analyzed per hypocotyl (total ≥15 cells per condition). All measurements used complete cell boundaries and identical image processing parameters. For *in vivo* HPTS calibration, hypocotyls were equilibrated in a measuring buffer (5 mM MES, 5 mM citric acid, 5M KOH, pH 2.5–7.5) for 30 min. The hypocotyls were then incubated in the same buffer supplemented with 5 mM HPTS for 20 min and subsequently mounted on a microscopy slide with the same buffer, covered with a coverslip, and imaged using the same confocal settings as described above. A calibration curve was generated and pH values were determined by fitting experimental data with a polynomial function.

##### ***In vivo* phosphorylation assay**

For the *in vivo* phosphorylation assay, 4-day-old seedlings were first subjected to darkness for 12 h, followed by 1 μM IAA treatment for 10 min and then harvested and flash frozen in liquid nitrogen. Samples were ground into a fine powder in liquid nitrogen and suspended in 2×SDS loading buffer (8% SDS, 100 mM Tris-HCl (pH 6.8), 20% Glycerol, 0.02% BPB) supplemented with protease inhibitor cocktail (A32955, Thermo Fisher Scientific, China). The homogenates were boiled at 99°C for 3 min and then centrifuged at 12,000 g for 10 min at room temperature. Equal amounts of supernatant were loaded onto 4%–12% (w/v) SDS–PAGE gels to assess the levels of AHA2 (anti-AHA2 1:1000 dilution, PY231, Beijing QiWei YiCheng Tech Co., Ltd) and phosphorylation at Thr947 (anti-pThr947, 1:1500 dilution, ZW075213, Abmart, China). Goat anti-rabbit IgG and goat anti-mouse IgG (1:5000 dilution) were used as secondary antibodies. The signals were detected using the Tanon-2500R imaging system (Tanon, China).

##### **Quantification and statistical analysis**

All experiments were independently repeated at least three times. Data are presented as mean ± SD. Statistical analyses were performed using GraphPad Prism 8 software. In all graphs, asterisks indicate statistical significance (n.s., not significant, *$P < 0.05$; **$P < 0.01$; ***$P < 0.001$; ****$P < 0.0001$) tested by Student's *t*-test (for two groups), and lowercase/uppercase letters indicate statistical significance determined between multiple groups by one/two-way ANOVA at $P < 0.05$ (lowercase) or $P < 0.01$ (uppercase). The intensity of protein bands in Western blotting was quantified using ImageJ software.

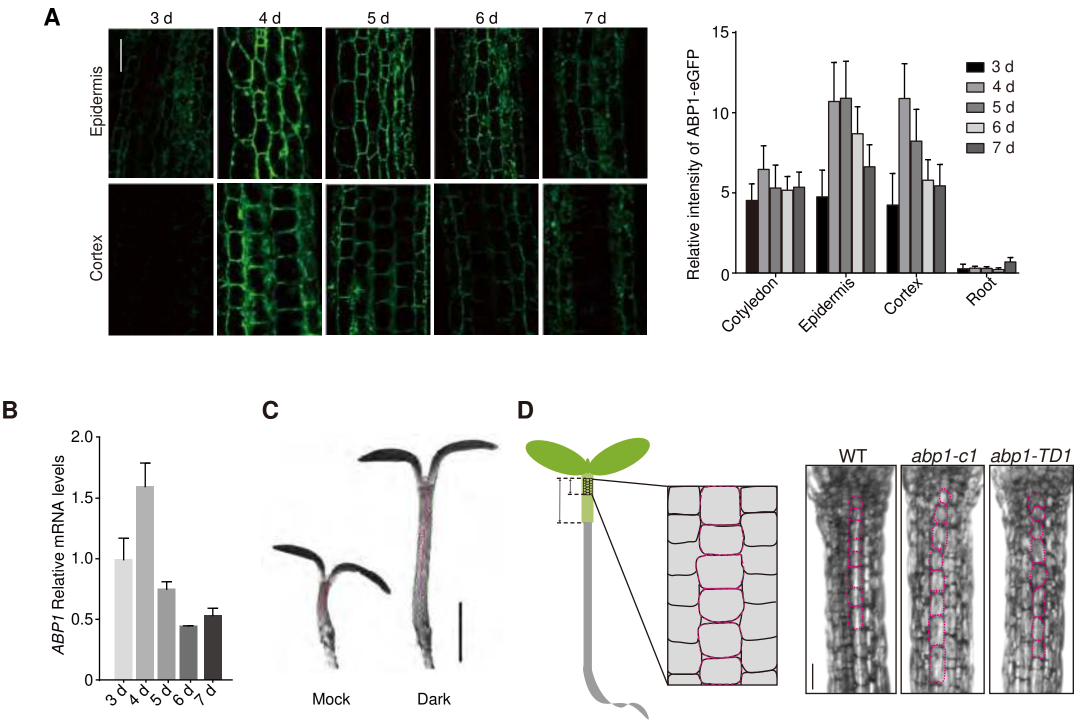


**Figure S1. Auxin regulates hypocotyl elongation under dark condition.**

(A) Confocal microscopy imaging showing the expression of ABP1 in the hypocotyl of *pABP1::ABP1-eGFP* transgenic plants from day 3 to 7. Scale bar, 50 µm. Time-course analysis of the fluorescence intensity of ABP1-eGFP in cotyledon, hypocotyl epidermis, hypocotyl cortex and root. $n \geq 14$ seedlings.

(B) RT-qPCR analysis of *ABP1* expression in aboveground part of seedlings from day 3 to 7.

(C) Representative images of hypocotyl cell segment under double-point tracking. Red lines indicate the tracking region. Scale bar, 1 mm.

(D) Schematic representation of the hypocotyl cell region examined during this study. Phenotype of hypocotyl cell areas in 4-day-old WT, *abp1-c1* and *abp1-TD1* seedlings subjected to darkness for 30 h. Scale bar, 100 µm.

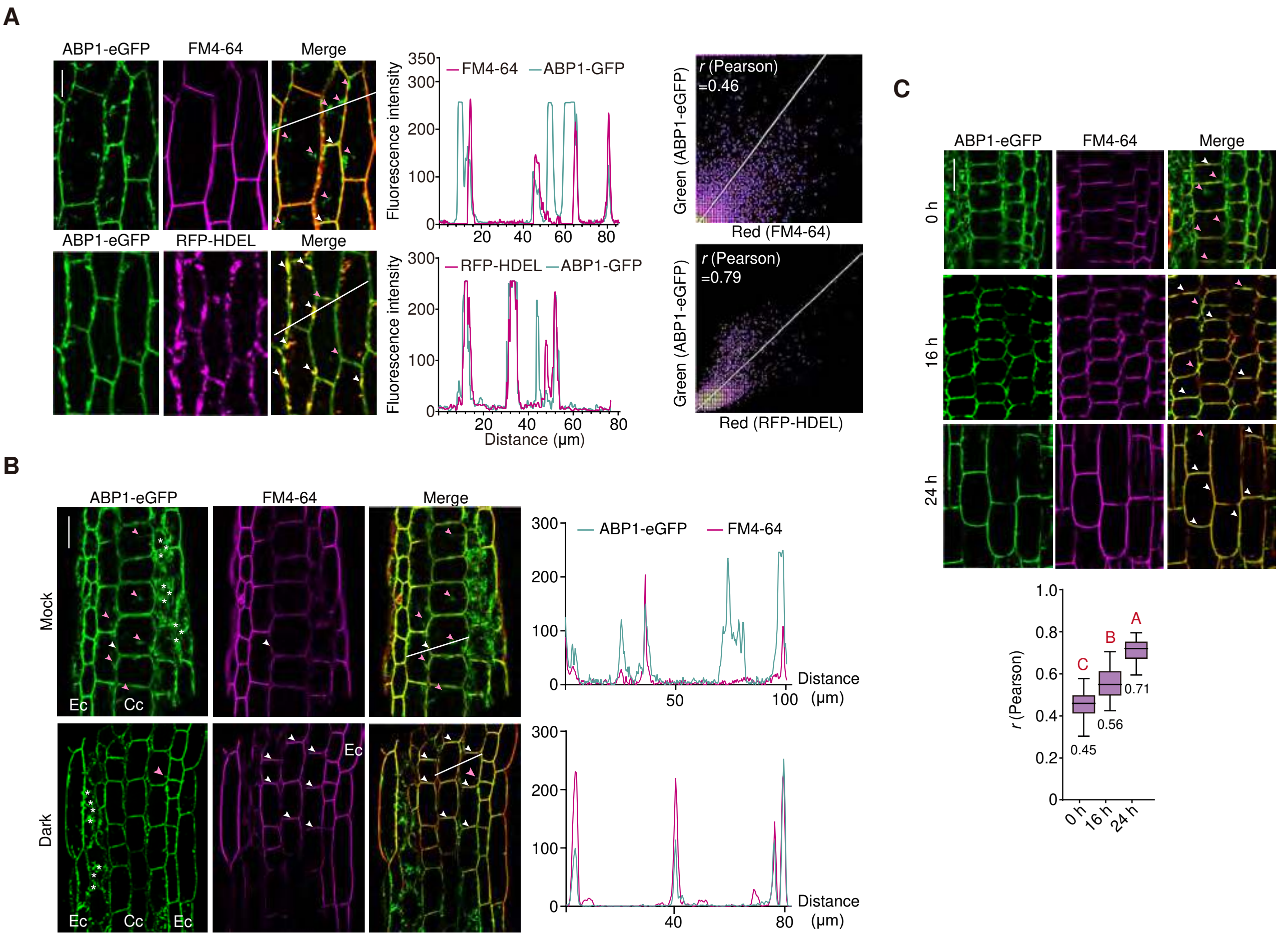


**Figure S2. Dark treatment induces the trafficking of ABP1.**

(A) Confocal microscopy imaging (left), fluorescence plot profiles (middle) and Scatterplot analysis (right) of ABP1-eGFP in hypocotyl epidermal cells of 4-day-old seedlings. FM4-64 staining indicates the PM. RFP-HDEL indicates the ER. (*r*), Pearson's colocalization coefficient. White arrowheads indicate the colocalization signal. Pink arrowheads indicate the non-colocalization signal. White lines are used for fluorescence intensity measurement. Scale bar, 20 μm.

(B) Confocal microscopy imaging and fluorescence plot profiling of ABP1-eGFP in hypocotyl cortical cells of seedlings subjected to dark treatment for 18 h. Ec, epidermal cell; Cc, cortex cell. stars lable the unfocused areas. White line is used for fluorescence intensity measurement; FM4-64 labels the PM; white arrowheads indicate the colocalized signal; pink arrowheads indicate the non-colocalized signal. Scale bar, 50 μm.

(C) Confocal microscopy imaging of ABP1-eGFP in hypocotyl cortical cells of seedlings subjected to dark treatment for 16 or 24 h. Pearson's coefficient showing the colocalization of ABP1-eGFP and FM4-64-stained PM. (*r*), Pearson's colocalization coefficient. White arrowheads indicate the colocalized signal. Pink arrowheads indicate the non-colocalized signal. $n \geq 15$ cells from 3 hypocotyls. Scale bar, 40 μm. Data are presented as mean ± SD. Significant differences were determined using one-way ANOVA. Different uppercase letters indicate significant differences at $P < 0.01$.

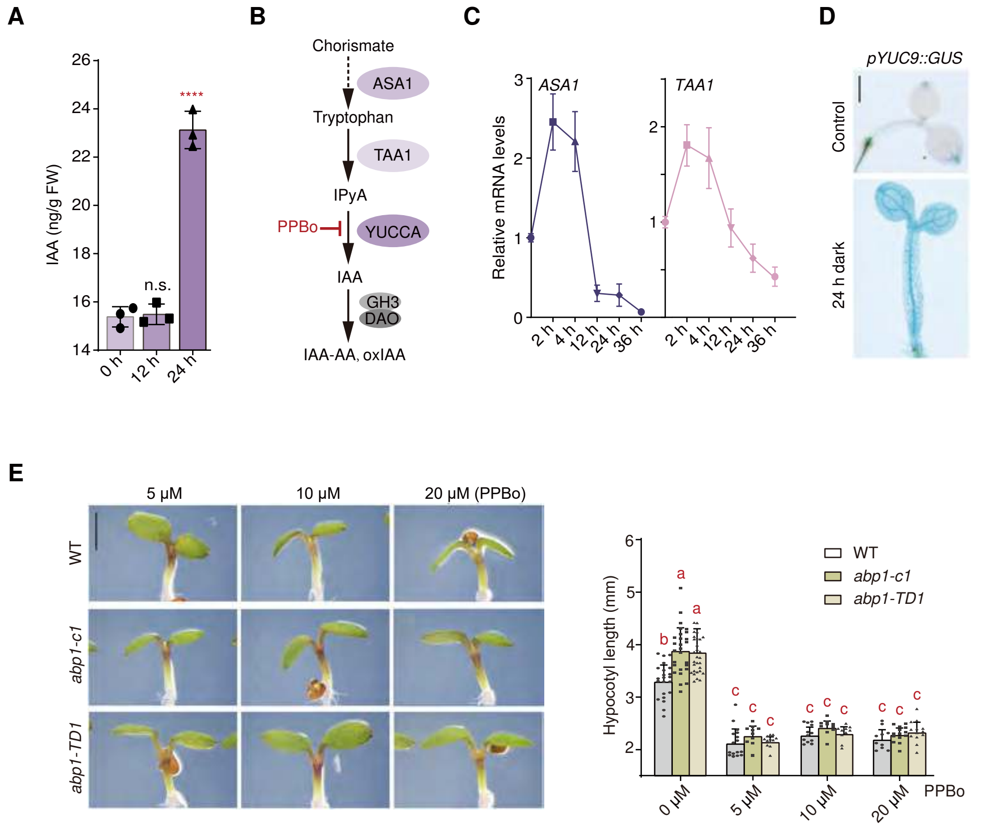


**Figure S3. Dark treatment induces auxin accumulation mediating hypocotyl elongation.**

(A) Free IAA contents in aboveground tissue of 4-day-old seedlings subjected to darkness for 12 or 24 h.

(B) Schematic representation of auxin biosynthesis pathway involving ASA1, TAA1 and YUCCA enzymes, with YUCCA activity inhibited by PPBo.

(C) RT-qPCR analysis of *ASA1* and *TAA1* in aboveground part of 4-day-old seedlings subjected to darkness for 0, 2, 4, 12, 24 or 36 h. Transcript levels were expressed as fold change relative to 0 h (set to 1).

(D) GUS staining in 4-day-old *pYUC9::GUS* transgenic seedlings under normal light conditions (top) and after 24 h dark treatment (bottom). Scale bar, 1 mm.

(E) Phenotype and quantification of hypocotyl length of 4-day-old light-grown WT, *abp1-c1* and *abp1-TD1* seedlings followed by 36 h co-treatment of darkness and auxin inhibitor targeting *YUCCA*s, PPBo. Scale bar, 2 mm. $n \geq 10$ hypocotyls.

In (A, E), data are presented as mean ± SD. Significant differences were determined using Student's *t*-test (A). ****$P < 0.0001$; n.s., not significant. Significant differences were determined using two-way ANOVA (E). Different lowercase letters indicate significant differences at $P < 0.05$.

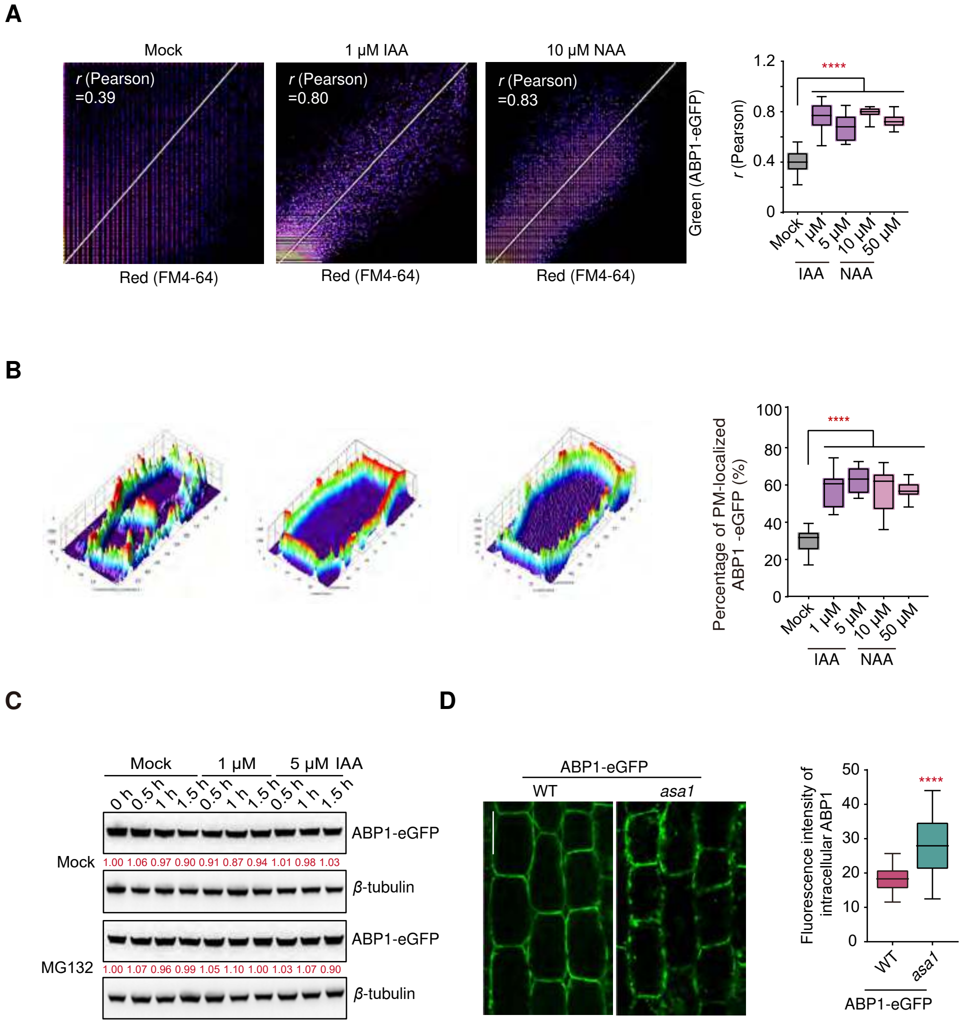


**Figure S4. IAA promotes the transport of ABP1 from the endoplasmic reticulum to the apoplast.**

(A) Scatterplot analysis and pearson's coefficient of ABP1-eGFP and FM4-64 colocalization after IAA or NAA treatment. $n \geq 13$ cells from 3–4 hypocotyls. ($r$), Pearson's colocalization coefficient.

(B) 3D surface plots and the percentage of PM-localized ABP1-eGFP in hypocotyl epidermal cells in 4-day-old seedlings treated with/without IAA or NAA for 1 h. $n \geq 15$ cells from 3–4 hypocotyls.

(C) Western blot showing the total protein levels of ABP1-eGFP in 4-day-old seedlings after IAA treatment at indicated period, with MG132 treatment as control. Numbers in red indicate protein amount of ABP1-eGFP after IAA treatment at each time point relative to non-treatment sample.

(D) Confocal microscopy imaging of ABP1-eGFP and fluorescence intensity of intracellular ABP1 in hypocotyl cortical cells of WT and *asa1* seedlings subjected to darkness for 18 h. Scale bar, 50 μm.

In (A, B, D), data are presented as mean ± SD. Significant differences were determined using Student's *t*-test (A, B, D). ****$P < 0.0001$.

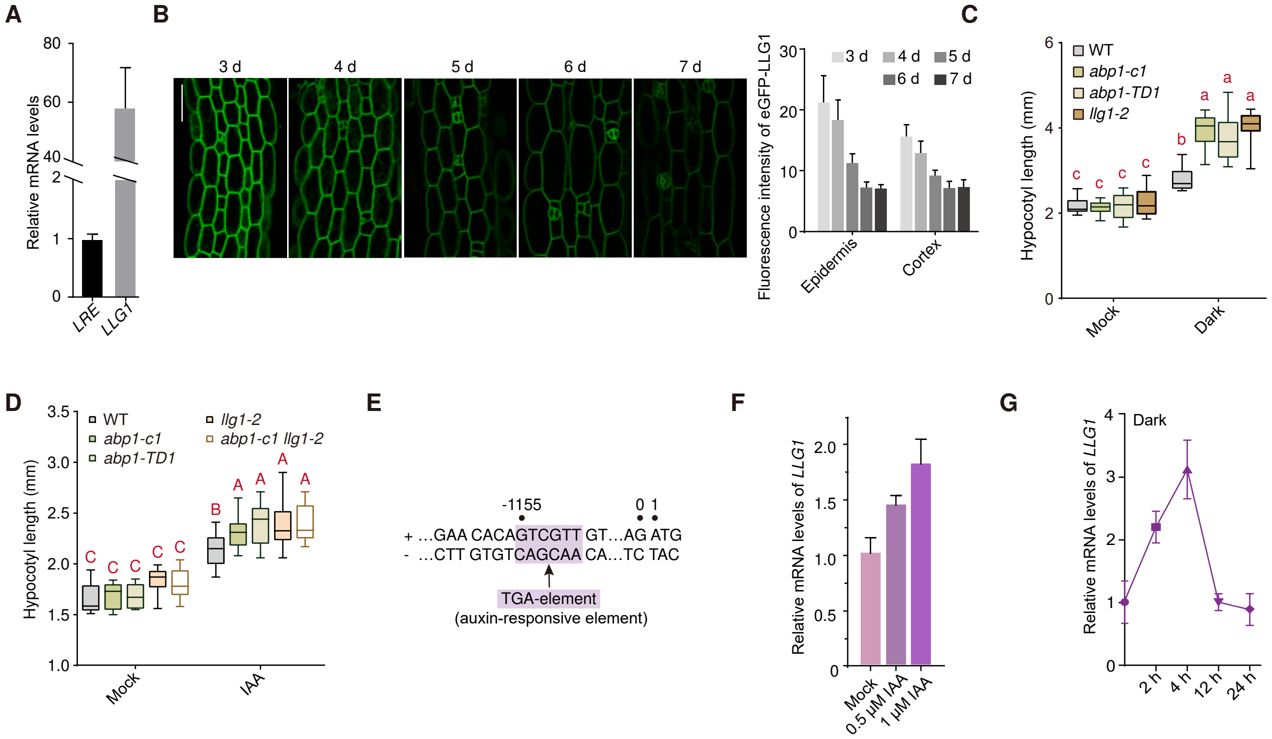


**Figure S5. LLG1 promotes hypocotyl elongation via auxin signaling.**

(A) RT-qPCR analysis of *LRE* and *LLG1* expression in the hypocotyl of 4-day-old seedlings.

(B) Confocal imaging showing the expression of LLG1 in hypocotyl cortical cells of *pLLG1:sp-eGFP-LLG1* transgenic plants from day 3 to 7. Scale bar, 50 μm. $n \geq 22$ seedlings. Time-course analysis of the fluorescence intensity of eGFP-LLG1 in hypocotyl epidermis and hypocotyl cortex from day 3 to 7.

(C) Quantification of hypocotyl length in 4-day-old WT, *abp1-c1*, *abp1-TD1 and llg1-2* seedlings subjected to dark treatment for 36 h. $n \geq 10$ hypocotyls.

(D) Quantification of hypocotyl length in WT, *abp1-c1*, *abp1-TD1*, *llg1-2* and *abp1-c1 llg1-2* seedlings under mock or IAA treatment for 12 h. $n \geq 9$ hypocotyls.

(E) Prediction in the *LLG1* promoter region showing an auxin-responsive element (TGA element).

(F) RT-qPCR analysis of *LLG1* expression in 4-day-old seedlings after IAA treatment for 0.5 h.

(G) RT-qPCR analysis of *LLG1* in aboveground tissues of 4-day-old seedlings subjected to darkness for 0–24 h. Transcription levels were indicated as fold change relative to 0 h (set to 1).

In (C, D), data are presented as mean ± SD. Significant differences were determined using two-way ANOVA. Different uppercase letters indicate significant differences at $P < 0.01$. Different lowercase letters indicate significant differences at $P < 0.05$.

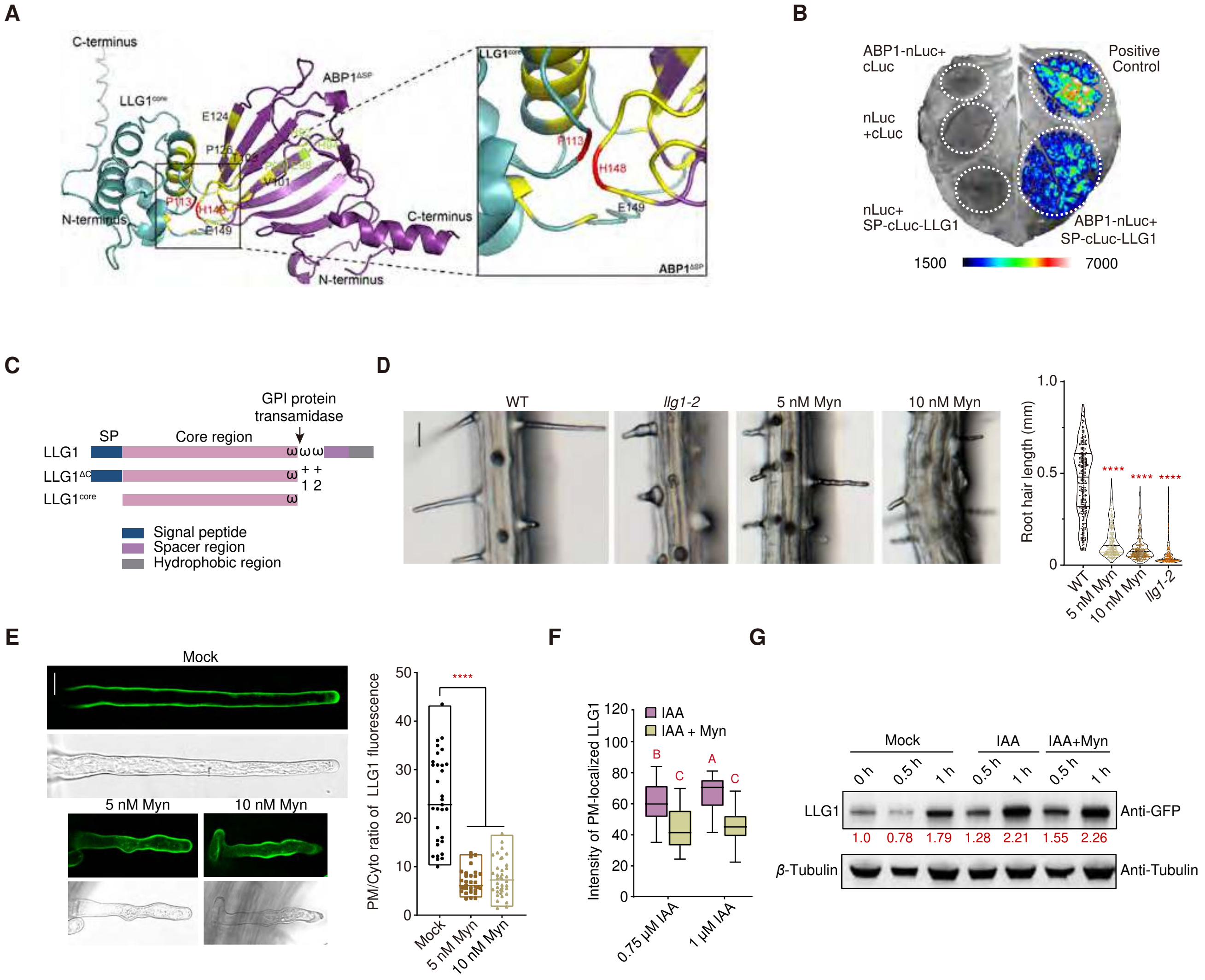


**Figure S6. ABP1 and LLG1 exhibit direct physical interaction and Myriocin blocks the LLG1-mediated transport.**

(A) AlphaFold3 protein interaction analysis of $ABP1^{\Delta SP}$ and $LLG1^{core}$. Yellow highlights the predicted $ABP1^{\Delta SP}$–$LLG1^{core}$ interaction interface. The right panel shows an enlarged view of the interface, with $LLG1^{core}$ shown in blue and $ABP1^{\Delta SP}$ shown in purple. The yellow region indicates the interaction interface and marks the 6 amino acid residues of $ABP1^{\Delta SP}$ involved in the predicted interaction. The green region highlights four key amino acid residues constituting the NAA-binding pocket.

(B) Luciferase complementation assay showing the interaction between full-length ABP1 and full-length LLG1. SP, signal peptide.

(C) The *LLG1* gene model, illustrating the predicted protein features, including the N-terminal signal peptide and the C-terminal GPI-anchor processing sites (ω-sites).

(D) Phenotype and quantification of root hair length in 3.5-day-old WT (with/without Myn treatment) and *llg1-2* (untreated) seedlings. Scale bar, 50 μm. $n \geq 149$ root hairs from 10 roots.

(E) Phenotype and quantification of PM/cytoplasm LLG1 fluorescence signal ratio of root hair in 3.5-day-old *pLLG1:sp-eGFP-LLG1* transgenic seedlings with/without Myn treatment. Scale bar, 20 μm.

(F) Quantification of fluorescence intensity of PM-localized eGFP-LLG1 in hypocotyl epidermal cells of 4-day-old *pLLG1:sp-eGFP-LLG1* transgenic seedlings treated with IAA alone or together with 50nM Myn. $n \geq 12$ cells from 3–4 hypocotyls.

(G) Western blot showing the total protein levels of eGFP-LLG1 in 4-day-old seedlings treated with 1 μM IAA alone or 50nM Myn co-treatment. Numbers in red indicate protein amount of eGFP-LLG1 after treatment at each time point relative to non-treatment sample.

In (D–F), data are presented as mean ± SD. Significant differences were determined using Student's *t*-test (D, E). ****$P < 0.0001$. Significant differences were determined using two-way ANOVA (F). Different uppercase letters indicate significant differences at $P < 0.01$.

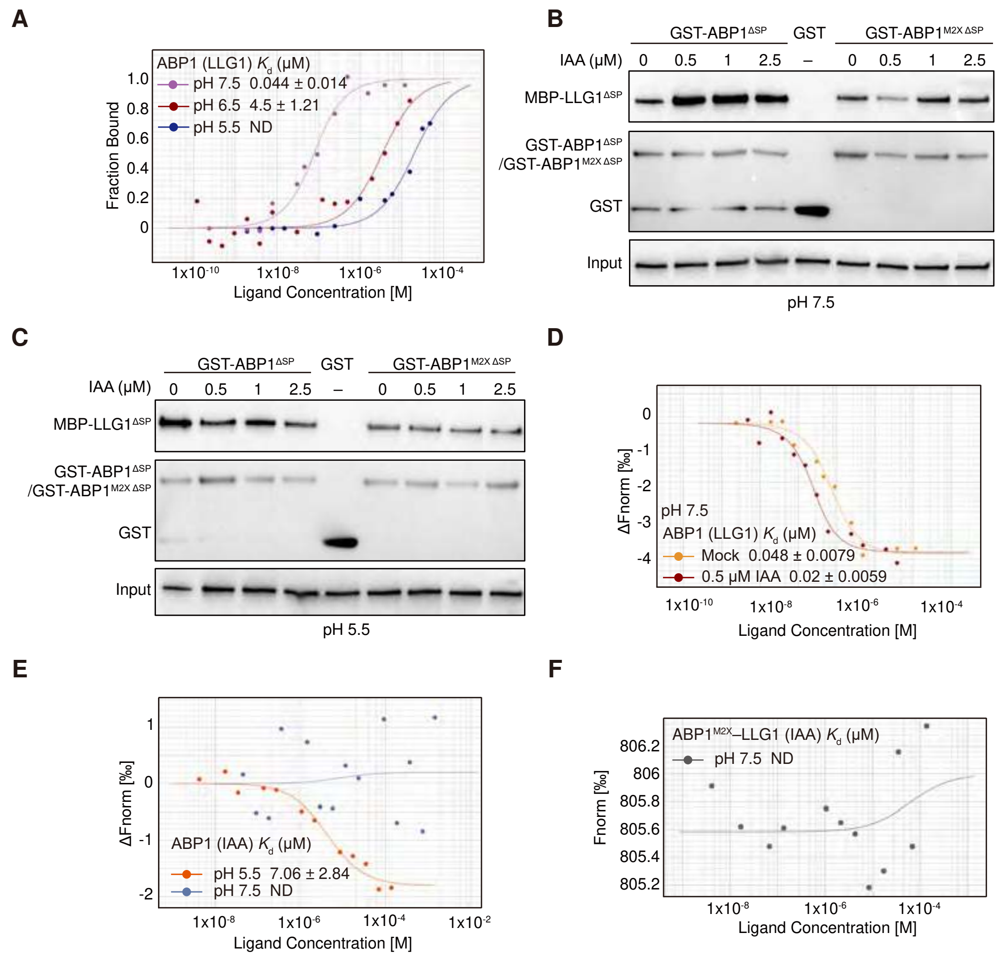


**Figure S7. Auxin and pH affect the interaction between ABP1 and LLG1.**

(A) MST analysis of the ABP1–LLG1 interaction under pH 5.5, pH 6.5 and pH 7.5 conditions.

(B,C) Pull-down assay showing $ABP1^{\Delta SP}$–$LLG1^{\Delta SP}$ or $ABP1^{M2X\ \Delta SP}$–$LLG1^{\Delta SP}$ interactions with various concentrations of IAA treatment under pH 7.5 or pH 5.5.

(D) MST analysis of the effect of IAA on the ABP1–LLG1 interaction under pH 7.5.

(E) MST analysis of IAA binding to ABP1 under pH 5.5 and 7.5.

(F) MST analysis of the IAA binding affinity to $ABP1^{M2X}$–LLG1 under pH 7.5.

(A, D–F) $K_d$, dissociation constant. Datapoints indicate the difference in normalized fluorescence (%) and the curves show calculated fits. Data are presented as mean ± SD. $n$ = 3 (A,F)); ≥ 3 (D,E). ND, no measurable binding signal observed.

A

B

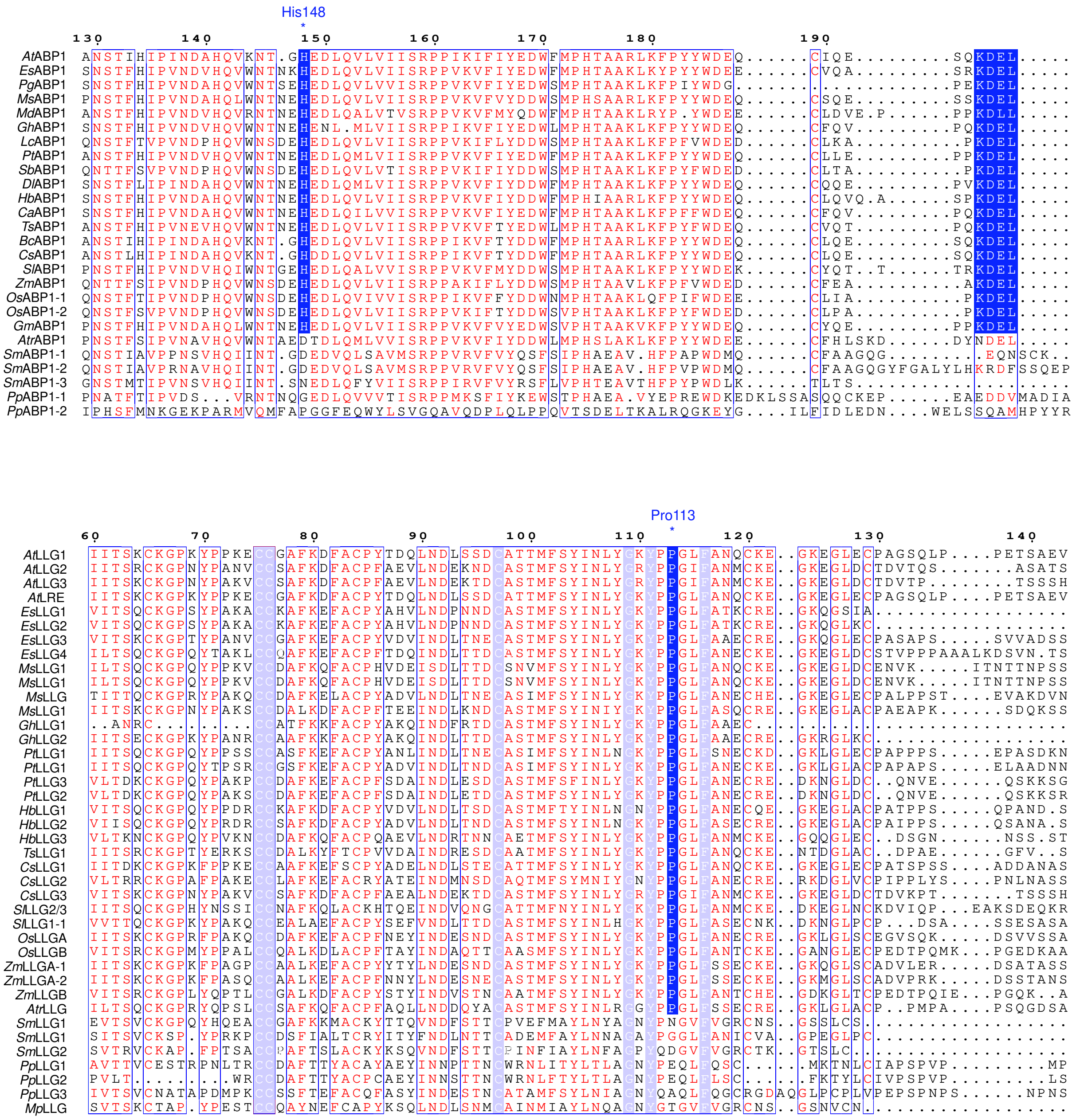

**Figure S8. Sequence alignment and conservation analysis of ABP1 and LLG1 proteins.**

(A) Sequence alignment of *At*ABP1 and its homologes across land plant species.

(B) Sequence alignment of *At*LLG1 and its homologes across land plant species.

*Eucalyptus spp (Es), Panax ginseng(Pg), Medicago sativa (Ms), Malus domestica (Md), Gossypium hirsutum (Gh), Leymus chinensis (Lc), Populus trichocarpa (Pt), Sorghum bicolor (Sb), Dimocarpus longan (Dl), Hevea brasiliensis (Hb), Corylus avellana (Ca), Turnera subulata (Ts), Brassica campestris (Bc), Camelina sativa (Cs), Arabidopsis thaliana (At), Zea mays (Zm), Oryza sativa (Os), Glycine max (Gm), Amborella trichopoda (Atr), Selaginella moellendorffii (Sm), Physcomitrium patens (Pp).*

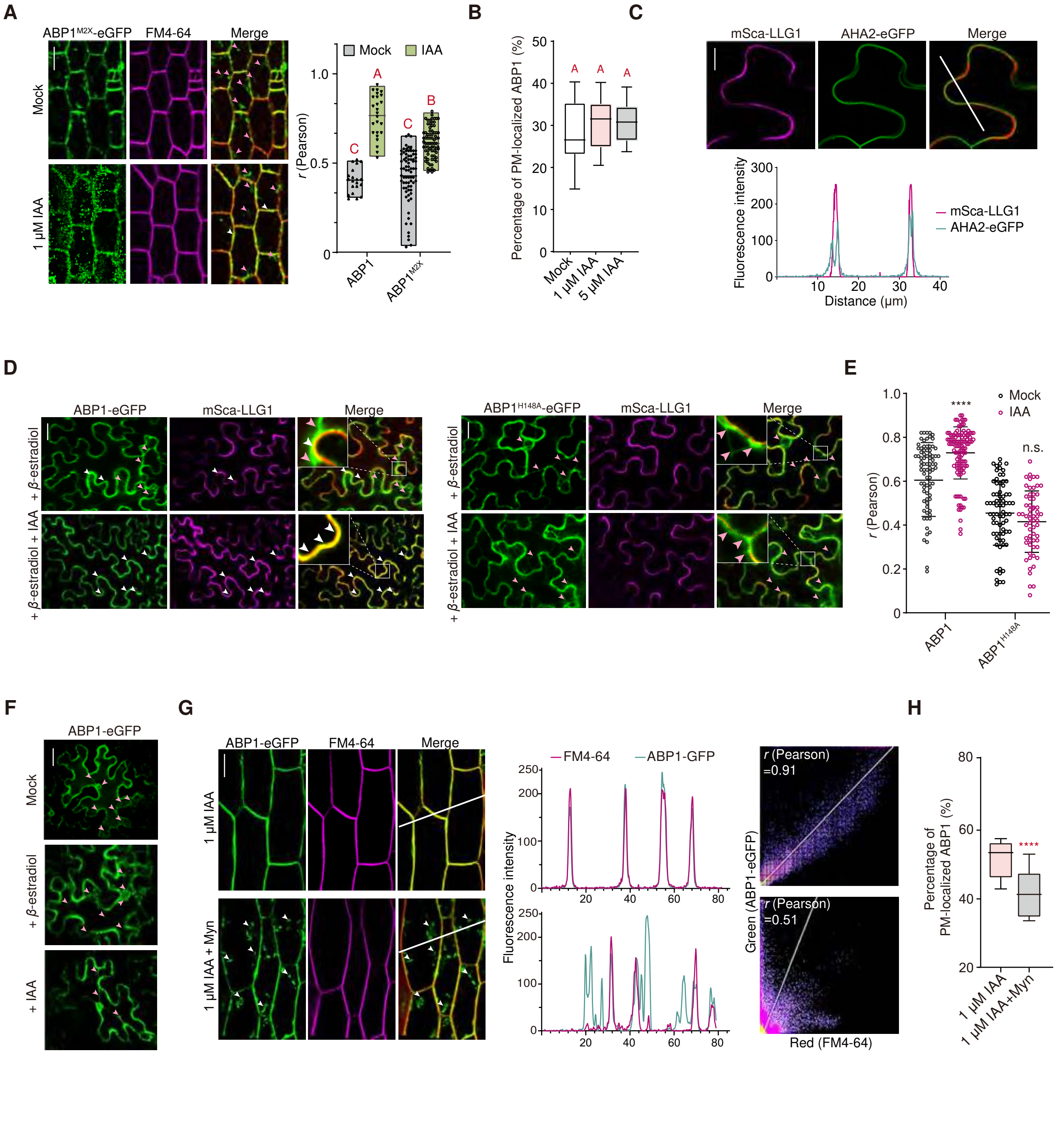

**Figure S9. LLG1 chaperones ABP1 transportation through GPI-AP pathway.**

(A) Fluorescence imaging and pearson's coefficient of hypocotyl epidermal cells in 4-day-old *pABP1:ABP1$^{M2X}$-eGFP/abp1-c1* seedlings treated with/without IAA for 1 h. White arrowheads indicate the colocalization signal. Pink arrowheads indicate the non-colocalization signal. Scale bar, 40 μm.

(B) The percentage of ABP1-eGFP localized to PM showing the colocalization of FM4-64 and ABP1-eGFP in *llg1-1* background after IAA treatment for 1 h. $n \geq 42$ cells from 15 hypocotyls.

(C) Confocal microscopy imaging and fluorescence plot profiles of mSca-LLG1 and PM marker AHA2-eGFP. White line indicates the region for fluorescence intensity measurement. Scale bar, 10 μm.

(D) Fluorescence imaging and pearson's coefficient of ABP1 (or ABP1$^{H148A}$) and LLG1 in tobacco leaves. *pABP1:ABP1-eGFP* and *pER8:mSca-LLG1* were transformed to *N. benthamiana* leaves and treated with *β*-estradiol for 24 h and 1 μM IAA treatment for 35 min. Pink arrowheads indicate the ER regions. White arrowheads indicate the regions of colocalization. Scale bar, 20 μm.

(E) Pearson's coefficient of ABP1 (or ABP1$^{H148A}$) and LLG1 in tobacco leaves. *pABP1:ABP1-eGFP* and *pER8:mSca-LLG1* were transformed to *N. benthamiana* leaves and treated with *β*-estradiol for 24 h and 1 μM IAA treatment for 35 min. $n \geq 61$ cells.

(F) Fluorescence imaging showing the expression of ABP1-eGFP alone under mock condition and treated with *β*-estradiol for 24 h or 1 μM IAA for 35 min in *N. benthamiana* leaves. Pink arrowheads indicate the ER regions. Scale bar, 20 μm.

(G) Confocal microscopy imaging (left), fluorescence plot profiles (middle) and scatterplot analysis with Pearson's colocalization coefficient (right) of ABP1-eGFP in hypocotyl epidermal cells of 4-day-old seedlings treated with IAA alone or 50nM Myn co-treatment for 1 h. White lines are used for fluorescence intensity measurement. White arrowheads indicate the punctate signals in ER. Scale bar, 20 μm.

(H) The percentage of ABP1-eGFP localized to PM relative to total protein in 4-day-old seedlings treated with IAA alone or 50 nM Myn co-treatment. $n = 5$ hypocotyls.

In (A, B, E, H), data are presented as mean ± SD. Significant differences were determined using two-way ANOVA (A). Significant differences were determined using one-way ANOVA (B) or Student's *t*-test (E, H). Different uppercase letters indicate significant differences at $P < 0.01$. n.s., not significant. ****$P < 0.0001$.

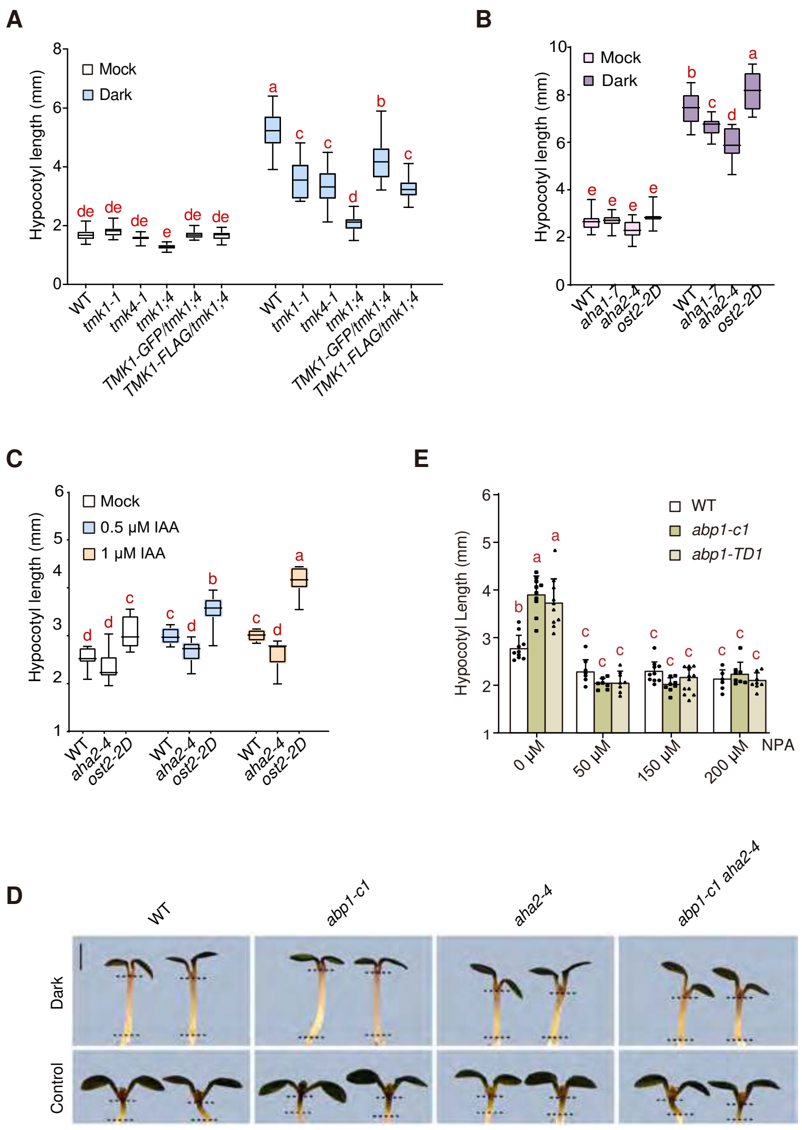


**Figure S10. AHA1/2 regulate hypocotyl elongation under dark and auxin treatment.**

(A) Quantification of hypocotyl length in *tmk1-1, tmk4-1, tmk1-1 tmk4-1, TMK1-GFP/tmk1 tmk4 and TMK1-FLAG/tmk1 tmk4* seedlings subjected to darkness treatment for 30 h. $n \geq 16$ seedlings.

(B) Quantification of hypocotyl length in *aha1-7, aha2-4* and *ost2-2D* seedlings subjected to darkness treatment for 30 h. $n \geq 16$ seedlings.

(C) Quantification of hypocotyl length in *aha2-4* and *ost2-2D* seedlings treated with/without IAA for 30 h. $n \geq 7$ seedlings.

(D) Phenotype of hypocotyl length in WT, *abp1-c1*, *aha2-4* and *abp1-c1 aha2-4* seedlings subjected to darkness for 30 h. $n \geq 23$ seedlings. Scale bar, 2 mm.

(E) Quantification of hypocotyl length in 4-day-old light-grown *abp1-c1* and *abp1-TD1* seedlings followed by 36 h co-treatment of dark and auxin inhibitor targeting *PIN*s, NPA. $n \geq 7$ hypocotyls.

In (A–C, E), data are presented as mean ± SD. Significant differences were determined using two-way ANOVA. Different lowercase letters indicate significant differences at $P < 0.05$.

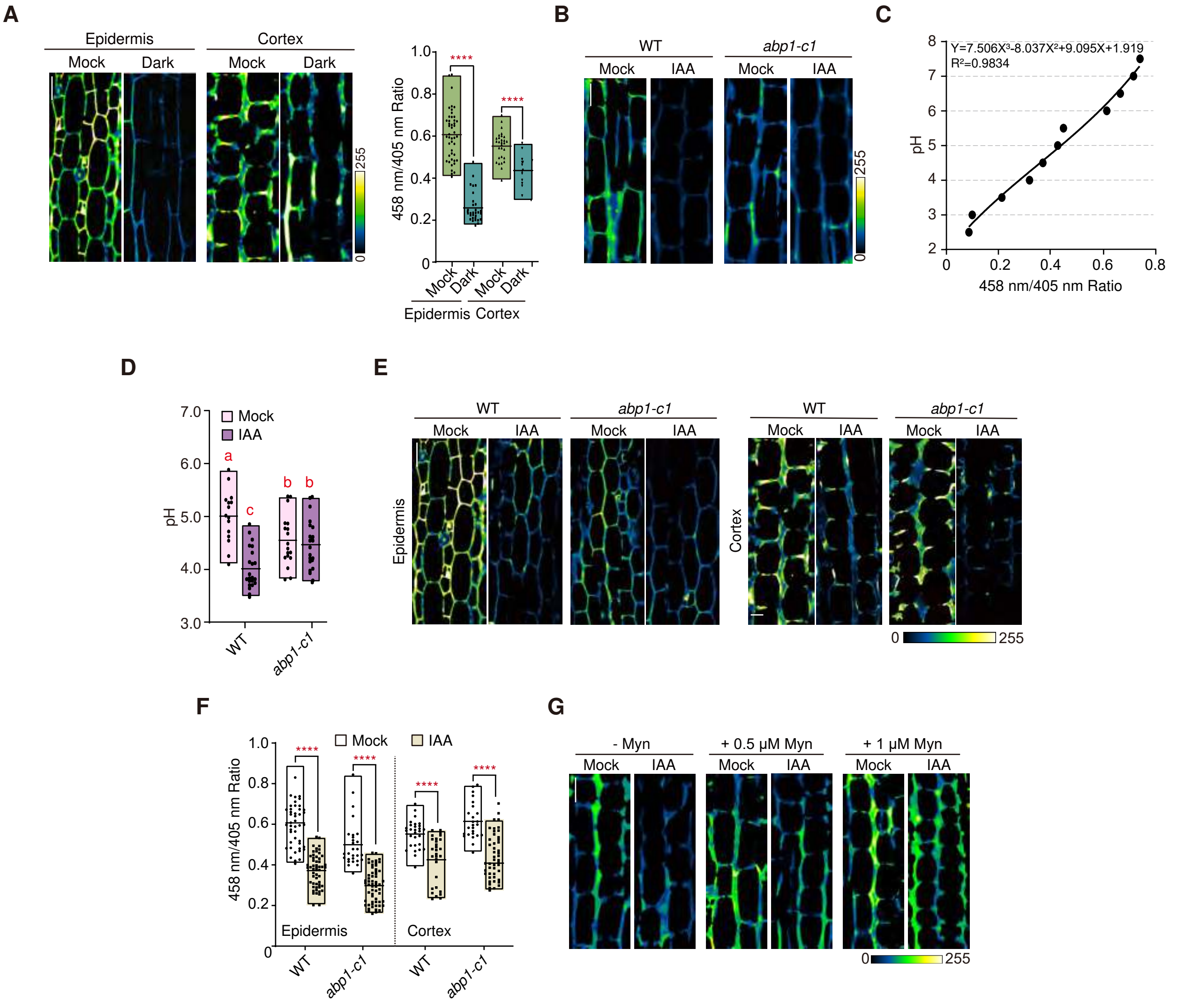


**Figure S11. ABP1 regulates dark-induced apoplast acidification in hypocotyl.**

(A) Confocal imaging and relative quantification of HPTS-stained apoplastic pH in hypocotyl epidermal and cortical cells in 4-day-old WT seedings subjected to darkness for 18 h. $n \geq 15$ cells. Scale bar, 40 μm.

(B) Confocal imaging of HPTS-stained apoplastic pH in hypocotyl cortex cells in 4-day-old WT and *abp1-c1* seedlings after dark treatment for 18 h following 1 μM IAA treatment for 15 min. Scale bar, 40 μm.

(C) *In vivo* calibration curve of apoplastic pH.

(D) Relative quantification of apoplastic pH in hypocotyl cortical cells of 4-day-old WT and *abp1-c1* seedlings subjected to darkness for 18 h, followed by 1 μM IAA treatment for 15 min. HPTS-stained pH values were calculated using a standard curve.

(E–F) Confocal imaging and relative quantification of HPTS-stained pH in the apoplast of hypocotyl epidermal and cortical cells in 4-day-old light-grown WT and *abp1-c1* seedlings treated with 1 μM IAA for 15 min. $n \geq 25$ cells. Scale bar, 40 μm.

(G) Confocal imaging of HPTS-stained pH in the apoplast of hypocotyl cortical cells in 3-day-old WT seedlings treated with Myn for 1 d followed by 18 h dark treatment and then 1 μM IAA treatment for 15 min. Scale bar, 40 μm.

In (A, D, F), data are presented as mean ± SD. Significant differences were determined using Student's *t*-test (A, F) or two-way ANOVA (D). ****$P < 0.0001$. Different lowercase letters indicate significant differences at $P < 0.05$.
