## Supplemental Table 1 for "Receptor trafficking couples intracellular auxin perception to rapid signaling"

**Table S1. List of primers used in this study.**

| Construct | Primer name | Sequence (5' to 3') | Purpose |
|---|---|---|---|
| *YUC3* | *F* | CGTTTGAATTAGGAGTTACG | qRT-PCR |
| | R | GGTATCCCATCATCGGAG | |
| *YUC8* | *F* | AATGCGGAGAGAGTGATGCC | |
| | R | TTCTCACGACCATGGAAGGC | |
| *YUC9* | F | GGCGATGTGTTTGGGTCAAC | |
| | R | CTCGACCACAACGAATGGGA | |
| *ASA1* | F | GAGTCAACGTTTTGAGCGGC | |
| | R | TTCTCTCCATCACCAACCTGC | |
| *TAA1* | F | GATGGGTGACAGGTGTACGG | |
| | R | CAGCGTTACCAACAACACCG | |
| *LLG1* | F | GCCTTCAAGGACTTTGCGTG | |
| | R | GTCCAGGCGGGTATTTTCCA | |
| *LRE* | F | CCCTCGTTTGTGGGAATGGA | |
| | R | TGAACGATTCCTGGACCTGC | |
| *Actin2* | *F* | CGTTTGAATTAGGAGTTACG | |
| | R | GGTATCCCATCATCGGAG | |
| GST-ABP1 | BamHI-F | TtccaggggcccctgggatccATGGCTCCTTGTCCCATCAATGG | Pull-down |
| | SalI-R | GatgcggccgctcgagtcgacAAGCTCGTCTTTTTGTGATTCTT | |
| | S106A-R | Tgcgccctttaggacaacaaa | |
| | S106A-F | TtgttgtcctaaagggcgcAGGTACTCTGTATCTCGCTGAAACAC | |
| | T146A-R | ctcatgaccgGCGTTTTTGACCTGATGAGCA | |
| | T146A-F | tcaaaaacgcCGGTCATGAGGACCTGCAGG | |
| | H148A-R | tcctcagCACCGGTGTTTTTGACCTGATG | |
| | H148A-F | aaaaacaccggtgCTGAGGACCTGCAGGTGTTGG | |
| pABP1: ABP1-eGFP | BamHI-F | gagctcggtacccggggatccAGTCTCGGAATACCAAGAACCTCG | Transgenic plant & agro-infiltration |
| | R | catTTTCTCGATGCTTCGACGAAC | |
| | F | gtcgaagcatcgagaaaATGATCGTACTTTCTGTTGGTTCCG | |
| | SalI-R | tgaaccgccaccgccgtcgacAAGCTCGTCTTTTTGTGATTCTTG | |
| | M2X-R | cacaggagACCCTGACAATTGGTGTCTCTGA | |
| | M2X-F | aattgtcagggtCTCCTGTGCCTGTGAAGAGGTT | |
| cLUC-LLG1 | SP-F | gagaacacgggggacgagctcATGGAGCTCCTCTCTAGAGCTCTTT | Split-Luc |
| | SP-R | aaccggacatTGAAGAAGAGAAAGAAGAAAGAACTGA | |
| | cL-F | ctcttcttcaATGTCCGGTTATGTAAACAATCCG | |
| | cL-R | aatgaaactACCTCCGCCAGATGAGCC | |
| | LLG1-F | ctggcggaggtAGTTTCATTTCAGATGGGGTCTTC | |
| | LLG1-R | tgtagtccatttgttggatccGAACAACTTAACAAAAACCAAAAGAGC | |
| | ΔGPI-R | tgtagtccatttgttggatccCGAGGTAGTTGCTGCGTTTACCTCTG | |
| ABP1-nLuc | Kpn1-F | acgggggacgagctcggtaccATGATCGTACTTTCTGTTGGTTCCG | |
| | Sal1-R | cgcgtacgagatctggtcgacAAGCTCGTCTTTTTGTGATTCTT | |
| | ΔKDEL-nL-R | cgcgtacgagatctggtcgacTTGTGATTCTTGAATGCATTGCT | |

| | | | |
|---|---|---|---|
| FRET-ABP1-LLG1/COB | attB3-ABP1 | ggggacaactttgtataataaagttgtaATGATCGTACTTTCTGTTGGTTCCG | FRET-FLIM |
| | attB2-ABP1 | ggggaccactttgtacaagaaagctgggtAAGCTCGTCTTTTTGTGATTCTTG | |
| | attB1-LLG1 | ggggacaagtttgtacaaaaaagcaggcttaATGGAGCTCCTCTCTAGAGCTCTTT | |
| | attB4-LLG1 | ggggacaactttgtatagaaaagttgggtgGAACAACTTAACAAAAACCAAAAGA | |
| | attB1-COB | GgggacaagtttgtacaaaaaagcaggcttaATGGAGTCTTTCTTCTCCAGA | |
| | attB4-COB | GgggacaactttgtatagaaaagttgggtgGGCAGAGAAGAAGAAAAAGAC | |
| pER8-mScarlet-LLG1 | SP-F | ctagtcgactctagcctcgagATGGAGCTCCTCTCTAGAGCTCTTT | agro-infiltration |
| | SP-R | cccttgctcaccatTGAAGAAGAGAAAGAAGAAAGAACTGA | |
| | mScarlet-F | tcttcaATGGTGAGCAAGGGCGAGG | |
| | mScarlet-R | tgagccacctccCTTGTACAGCTCGTCCATGCC | |
| | LLG1-F | tgtacaagGGAGGTGGCTCATCTGGCG | |
| | LLG1-R | gggaggcctggatcgactagtTCACTTGTCATCGTCATCCTTGTAA | |
| | LLG1$^{\Delta GPI}$-R | gggaggcctggatcgactagtTCAAGTTGCTGCGTTTACCTCTG | |
| *abp1-TD1* | LP | ATGATCGTACTTTCTGTTGGTTCC | Genotyping |
| | RP | TTAAAGCTCGTCTTTTTGTGATTCT | |
| | pSKTAIL-L | ATACGACGGATCGTAATTTGTCG | |
| *abp1-c1* | genomic-F | TTCGTCGAAGCATCGAGAAA | |
| | genomic-R | TTTTAGCCTCGGACTTTGCT | |
| ABP1$^{M2X}$ | M2X-R | cacaggagACCCTGACAATTGGTGTCTCTGA | |
| | M2X-F | aattgtcagggtCTCCTGTGCCTGTGAAGAGGTT | |
| *llg1-2* | *llg1-2* LP | ATGGAGCTCCTCTCTAGAGCTC | |
| | *llg1-2* LB | GCTTCCTATTATATCTTCCCAAATTACCAATACA | |
| | *llg1-2* RP | TCAAGAGGATCATCATTAAGCCAC | |
| *aha1* | *aha1-7* LP | GCGTTGTAACTCTTGCAGTTTG | |
| | *aha1-7* RP | TCTTTCTTGGTTGTGAAAGCG | |
| | *aha1-6* LP | CGTCTCAACAAAAGTCTCTTTCA | |
| | *aha1-6* RP | CGAAAGATCAACCTCGTGAGT | |
| *aha2* | *aha2-4* LP | TTGAAAAGGCTGATGGATTTG | |
| | *aha2-4* RP | CTCCAGGACGTTCAACAAAAG | |
| *pAHA2: AHA2* | proF | CTATGACATGATTACGAATTCgcaacttttgaaaaagatgaatgg | Transgenic plant |
| | proR | ACTCGACATctctcaccactccttcactgagaa | |
| | CDS-F | agtggtgagagATGTCGAGTCTCGAAGATATCAAGAA | |
| | T881A-R | AACCGTGAAGTGCCCTTTGAGCAAGTGCCC | |
| | T881A-F | TCAAAGGGCACTTCACGGTTTACAGCCAAAAGA | |
| | T947A-R | AGGTCGACTCTAGAGGATCCTCACACAGCGTAGTGACTGGGAG | |
